# MoLst8 regulates autophagy and lipid homeostasis in *Magnaporthe oryzae*

**DOI:** 10.1101/2024.05.07.592880

**Authors:** Xingwei Cao, Lin Li, Jiandong Bao, Jiaoyu Wang, Xiaohong Liu, Xueming Zhu, Fucheng Lin

## Abstract

TOR, a widely conserved eukaryotic protein kinase, forms TORC1 and TORC2 to regulate diverse cell signaling. TORC1 controls protein synthesis, cell cycle, and autophagy, whereas TORC2 manages cell polarity, cytoskeleton, and membrane structure. Our previous research found that MoVast2, along with MoVast1, regulates TOR in rice blast fungus *Magnaporthe oryzae*, maintaining lipid and autophagy balance. Lst8, a key TOR complex component in yeast and mammalian cells. However, the precise role of MoLst8 in *M. oryzae* is still unclear. In this study, we obtained the Δ*Molst8* mutant through high-through gene knockout strategies. The results showed that loss of *MoLST8* leading to a series of defects, such as growth and sporulation reduction, abnormal conidia, and loss of virulence. In addition, this mutant is highly sensitive to rapamycin, leading to growth arrest and autophagy impairment, indicated that MoLst8 positively regulates TORC1 for cellular growth, metabolism, and autophagy. Lipidomics analysis in the mutant revealed lipid metabolism dysregulation, sphingolipid reduction, disrupting membrane tension and homeostasis, suggested that TORC2 mediated lipid regulation is disordered in Δ*Molst8* mutant. Additionally, the study explored TOR-MAPK crosstalk, finding that the mutant shows heightened cell wall stress sensitivity but fails to restore integrity despite MAPK activation. These findings offer insights into MoLst8’s role in fungal pathogenesis, contributing to an understanding of fungal biology and disease control strategies.

## INTRODUCTION

In recent years, rice blast fungus has become rampant in temperate and tropical regions, causing substantial economic losses to crops [1, 2]. The significant decline in global rice production has emerged as a serious threat to food security, capturing the attention of nations worldwide [3, 4]. *Magnaporthe oryzae*, prevalent in 85 countries with diverse ecological settings, presents a significant menace as it primarily targets cultivated rice (*Oryza sativa*), a crucial staple sustaining above 50% of the world’s population [5]. In a conducive environment, its invasion can have devastating consequences for rice, reducing harvests up to 30%, and in severe cases, leaving rice grains unproductive [2, 6]. *M. oryzae* is declared to be the most globally destructive pathogen [5]. It has the capacity to afflict rice at their growth stages and invade various tissues, including leaf blades, stems, panicle necks, spikelet stalks and grains [2]. The infected lesions appear as elliptical or spindle-shaped spots with deeper edges, and a single spot can produce spores for more than 20 days, producing thousands of spores a night. Once they merge with the surrounding lesions, they cause the entire leaf to wither. Infections on the nodes result in stem breakage, and infections on the panicles lead to the complete necrosis of the panicle, preventing seed production. Infections at the leaf collar, where the leaf blade connects with the leaf sheath, result in the necrosis of the entire leaf. The conidia it produces consists of a clusters of three cells, resembling tear drops [7]. Utilizing the mucilage held at their tips, these conidia firmly attach to the surface of their host, initiating germination and the production of germ tubes [8]. The demise of these conidia acts as a catalyst for the formation of specialized structural appressoria. Subsequently, these appressoria undergo darkening and glycerol accumulation, resulting in their expansion and the creation of inward tension. This pressure aids their infiltrating of the host’s epidermal cells, enabling the invasive mycelium to establish colonization within the plant. Furthermore, they possess the capacity to influence the development of intercellular hyphae, enabling intercellular movement and, in turn, generating conidia that are dispersed in the air through airflow [9].

Autophagy is a cellular mechanism that involves the degradation and recycling of components within the cytoplasm, including organelles and macromolecules, in response to various stresses or developmental cues [10–13]. In recent years, autophagy was confirmed to contribute significantly to many pathogenic fungi. In rice blast fungus, autophagy-associated genes including *MoATG1*, *MoATG7*, *MoATG8*, *MoATG9* and *MoATG14*, have been documented to control the formation of the appressorium—a unique fungal structure utilized for infecting host plants. [14–16]. The development of appressoria is coupled with autophagy-induced the demise of conidial cells, supplying nourishment and energy necessary for the appressorial function. [17, 18]. In the wheat disease agent *Fusarium graminearum*, autophagy key proteins FgAtg1-10, FgAtg12, FgAtg14-18 were required for pathogenicity and deoxynivalenol (DON) toxin biosynthesis [28]. In *Harpophora oryzae*, an endophytic fungus, the removal of the autophagy gene *HoATG5* led to heightened vulnerability to agents that disrupt the cell wall, including SDS, CFW, and Congo red [26]. Recently, few autophagy regulate proteins were identified and confirmed to play vital roles in autophagy pathway. MoSnt2, a histone h3 deacetylation enzyme, oversees MoTOR-dependent autophagy and aids in the infection of plants by *M. oryzae* [6]. The novel vacuolar protein Imp1 is particularly important for maintaining membrane homeostasis and TOR-independent autophagy activation [19]. Nowadays, two new autophagy regulate proteins MoVast1 and MoVast2 were verified that they can control autophagy flux by mediating TOR activity and plasma membrane tension [24]. These results pointed towards the vital importance of autophagy in maintaining cellular homeostasis, regulating fungal cell death, and facilitating the development of infection structures. Understanding mechanisms of autophagy in *M. oryzae* could provide significant perspectives for formulating fresh tactics to curb rice blast disease [16].

Rapamycin target protein (TOR), a serine/threonine kinase conserved across eukaryotes, governs protein synthesis, cell cycle progression, mitosis, polarity, and cell homeostasis [29, 30]. Mammals possess two distinct mTOR complexes, mTORC1 and mTORC2. mTORC1 comprise of five components: mTOR, Raptor, mLST8, PRAS40, and DEPTOR. Its primary function is to govern cell growth and metabolism, and it is responsive to rapamycin. On the other hand, mTORC2 comprises six components, including mTOR, Rictor, mLST8, Deptor, mSIN1, and Protor1/2. mTORC2 chiefly governs cellular survival, growth, and the restructuring of the cytoskeleton [32]. Research shows mTOR’s involvement in essential cellular procedures, encompassing protein synthesis to autophagy. Its hyperactivation links to cancer, diabetes, and aging, occurring in 70% of human cancers [33]. In *Saccharomyces cerevisiae*, there exist two TOR kinases, namely Tor1 and Tor2. Among these, Tor1 forms the TORC1 complex by binding with Kog1, Tco89, and Lst8. It is primarily localized around the vacuolar or endosomal membranes and is responsible for regulating cell growth, nutrient absorption, and autophagy [34]. The TORC2 complex consists of Tor2, Avo1, Avo2, Avo3, Bit61, and Lst8. The TORC2 complex is pivotal in preserving the stability and integrity of the cell membrane [35]. When sphingolipids are reduced or the plasma membrane undergoes changes, the TORC2-Ypk1 signaling pathway is activated to promote sphingolipid synthesis, maintaining plasma membrane homeostasis, integrity, fluidity, signal transduction, and other functions [36, 37]. Research shows that myriocin-induced sphingolipid synthesis inhibition phosphorylates and inactivates Orm1/Orm2, boosting sphingolipid production. This is mediated by Ypk1, sensing sphingolipid levels and activating TORC2-Ypk1 signaling via Orm protein phosphorylation [38]. These researches suggested that TOR pathway plays key roles among eukaryotes. MAPK (Mitogen-activated protein kinase) cascades constitute conserved signaling systems in eukaryotes that range from cellular growth and differentiation to responses to various environmental stresses [39–41]. In fission yeast, Tor1/Tor2 sense nutrition, impacting cell cycle and sex differentiation. MAPK stress response pathway (SRP) oversees cellular size and mitotic processes, adapting to nutrition supply. TOR and MAPK, despite their differences, are interconnected for cell growth and division [42]. In *Schizosaccharomyces pombe*, it has revealed a intricate crosstalk existing between the TOR and MAPK signaling pathways. Ryh1, an activator of TORC2, can activate Pmk1, a core member of the MAPK pathway, while stress-activated Pmk1 can also inhibit Ryh1 signaling. This crosstalk involves both signal activation and inhibition, which is crucial for cellular adaptability and survival, especially during changes in phosphoinositide metabolism [43].

*Magnaporthe oryzae*’s rapamycin target protein (MoTOR) is a preserved serine/threonine kinase in eukaryotes that oversees cell growth, metabolism, and pathogenicity. In *S. cerevisiae* and mammalian cell, Lst8, as a component of TORC1 and TORC2 complexes, is the only protein in the TORC1 and TORC2 complex that co-exists. It binds to TOR kinase and provides support for the full catalytic activity of TOR, which act as cellular sensors for nutritional status [71, 72]. Studying Lst8 provides insights into how cells perceive their nutritional environment and transmit signals for growth regulation, balancing biosynthetic and degradative processes. The mutation in the *LST8-1* gene that will be significantly expressed in *Arabidopsis*, while not dying, will lead to stunted growth, subsequent delayed flowering, and an over-sensitivity too short to long day [73]. A recent study found that in *Chlamydomonas reinhardtii*, the mutation of *LST8* disrupts TORC1 signaling and cellular response to phosphorus starvation, leading to elevated triacylglycerol accumulation under both phosphorus-replete and phosphorus-limited conditions [74]. However, in plant pathogenic fungi, the constituents of the TORC2 complex and its role are predominantly unexplored. In our investigation, we pinpointed a postulated TOR subunit protein MoLst8 and obtained *MoLST8* knockout mutant. The Δ*Molst8* showed a series of defects, such as growth and sporulation reduction, abnormal conidia, and loss of virulence. In addition, this mutant is highly sensitive to rapamycin, leading to growth arrest and autophagy impairment. Lipidomics analysis in the mutant revealed lipid metabolism dysregulation, sphingolipid reduction, disrupting membrane tension and homeostasis. Additionally, we find that the mutant shows heightened cell wall stress sensitivity but fails to restore integrity despite MAPK activation. Our research provides comprehensive insights into the multifaceted role of MoLst8 in *M. oryzae*, revealing its involvement in TORC1 and TORC2 activities, growth, development, toxicity, autophagy regulation, sphingolipid homeostasis, membrane dynamics, and cellular responses to environmental stresses. The interconnection between TOR and MAPK pathways, mediated by MoLst8, emerges as a crucial regulatory axis orchestrating essential cellular processes for the survival and virulence of *M. oryzae*. These findings deepen our comprehension of fungal pathogenicity and open the path for further exploration of molecular mechanisms governing critical cellular processes in *M. oryzae*. Its multifunctionality positions Lst8 as a key regulatory node in eukaryotes, offering insights for understanding TOR signaling and potential avenues for developing therapeutic approaches for various diseases.

## RESULTS

### Identify the TOR subunit MoLst8 in *M. oryzae*

To explore TOR complexes in *M. oryzae*, we searched the whole genome of rice blast fungus and found some TOR complex consisted of proteins. The most important component of TOR complexes, the *MoLST8* cDNA, has a total length of 2.192 kb and consists of 4 exons, namely MGG_07284-E1 (258 bp), MGG_07284-E2 (22 bp), MGG_07284-E3 (496 bp), and MGG_07284-E4 (1075 bp), and 3 introns, namely intron 1-2 (127 bp), intron 2-3 (103 bp), and intron 3-4 (112 bp) (Fig. S1A). Therefore, a sequence of 317 amino acid residues was encoded, predicting a molecular weight of 35.10 kDa and a theoretical pI of 6.18. *MoLST8* gene produces MoLst8, which serves as a component of the protein target-rapamycin complex. Comparison of this protein with Lst8 proteins from fungal pathogens in the NCBI database showed a high level of conservation and homology, with a similarity of over 80% (Fig. S1B). We constructed a tree to visualize the relationships between the MoLst8 protein and a dataset of highly conserved pathogenic fungal proteins, which is presented in Fig. S2A. To predict the structure of the MoLst8 protein (XP_003715510.1), we used Swiss-Model for homology modeling. We derived an AlphaFold DB model template from the homologous protein encoded by the gene A0A4Q4XXX6_9PEZI in *Monosporascus sp* CRB-8-3, which highly matches the target sequence with a sequence similarity of 86.12%. We used the homologous protein A0A4Q4XXX6.1 as a template to construct a structural model, as shown in Fig. S2B.

### The characteristic of *MoLST8* deletion mutant in *M. oryzae*

In order to explore the function of the TORC2 complex in *M. oryzae*, we try to knock out these genes of TOR complex through the gene knockout approach outlined by Lu et al. [75]. After many rounds of screening, we obtained *MoLST8* detection mutant (Fig. S3). To test the impact of MoLst8 in cell growth and conidiation, we cultured Guy11 and Δ*Molst8* on complete media (CM) for a duration of 9 days at 25°C.. Some significant morphological differences were observed among Δ*Molst8* and Guy11. Firstly, the mycelium of Δ*Molst8* appeared white and folded (Figure 1A). Secondly, Δ*Molst8* exhibited slower growth and a significantly reduced conidiation rate compared to Guy11 (Figure 1B and 1C). Next, we studied the conidial morphology of Guy11, Δ*Molst8*, and Δ*Molst8-C*. In Δ*Molst8*, compared with Guy11 and Δ*Molst8-C*, the conidia failed to develop normally and exhibited abnormal morphology. When collecting conidia on CM plates, they were easily broken (Figure 1D and 1E). To test whether MoLst8 is involved in the virulence of *M. oryzae*, virulence tests were conducted on rice and barley, two distinct hosts. We added spore droplets (5 × 10^4^ spores/ml) of Δ*Molst8*, wild type, and Δ*Molst8-C* on barley and rice seedlings (CO-39) that were two weeks old. Seven days after inoculation, the Δ*Molst8* produced only small lesions, whereas the wild-type and Δ*Molst8-C* induced numerous, characteristic fused lesions, forming a clear contrast (Figure 1F and 1K). We conducted host penetration assays to further understand the impact ofΔ*Molst8* on disease progression in barley. Mycelia plugs of Guy11, Δ*Molst8*, and Δ*Molst8-C* were placed onto excised barley and rice leaves. After four days, Guy11 and Δ*Molst8-C* induced severe lesions, whereas the Δ*Molst8* mutant did not cause any lesions (Figure 1H and 1J). Subsequently, we conducted invasive hyphae (IH) assays to assess Guy11, Δ*Molst8*, and complemented strain infection on barley leaves. In Guy11 and the complemented strain, appressoria developed normally to form IH structures. The IH then colonized adjacent cells, often more than two cells. In contrast, approximately 30% of the Δ*Molst8* appressoria developed IH structures, and their ability to colonize adjacent cells was weak (Figure 1G and 1I). Overall, the Δ*Molst8* mutant exhibited significant growth retardation, reduced sporulation with abnormal morphology, and difficulty in forming IH structures to colonize adjacent host cells. The virulence was essentially lost. Our findings suggest that the *MoLST8* gene is pivotal for normal growth and development of *M. oryzae* mycelia, conidia, and appressoria as well as virulence.

**Figure 1.**
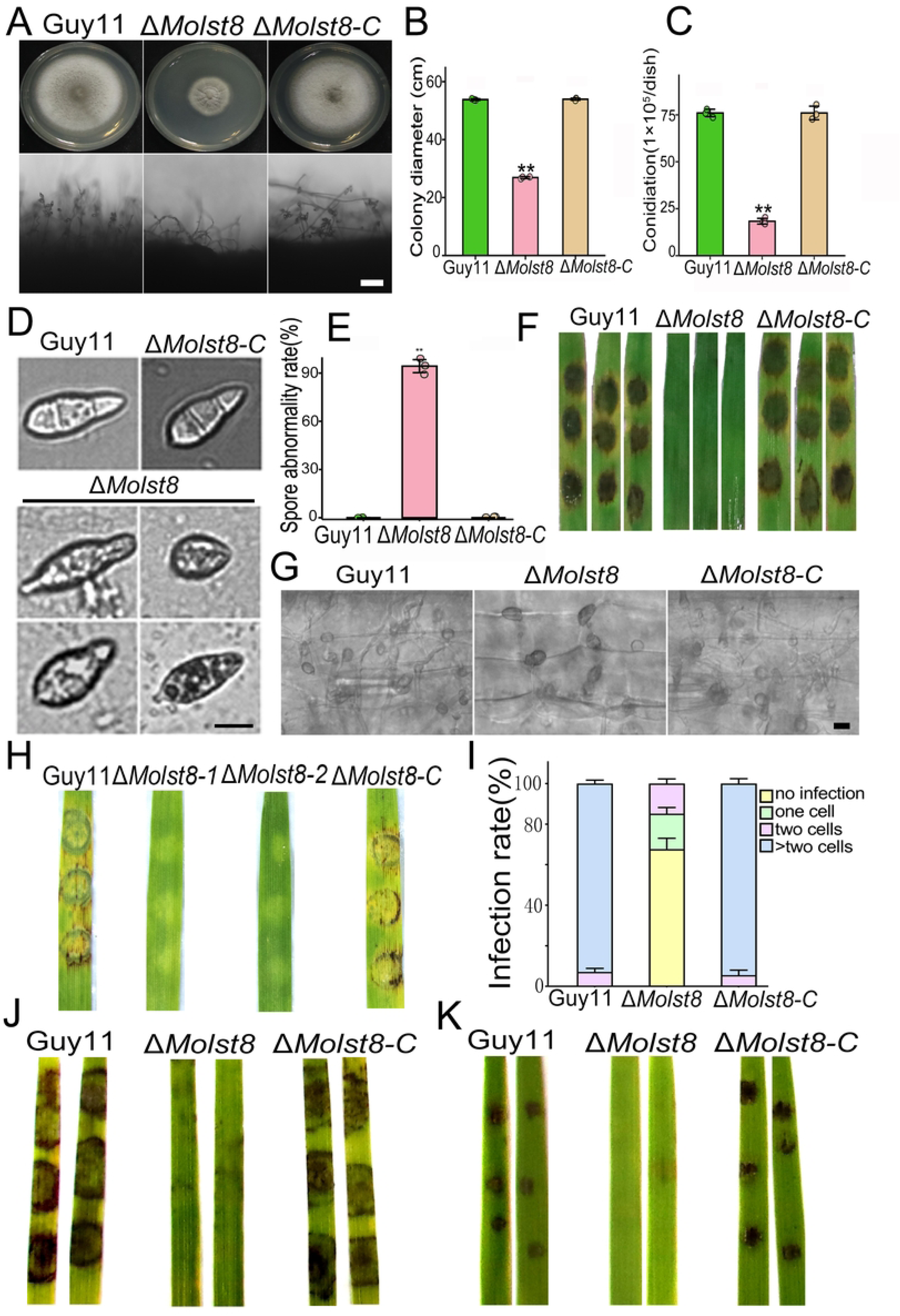
MoLst8 is involved in growth, development, and virulence. (A) The Colony morphology and conidiophores of Guy11, Δ*Molst8*, and the complemented strain, bar = 50 μm. (B) and (C) The colony growth rate and conidiation in Guy11, Δ*Molst8*, and the complemented strain. Data were examined with GraphPad Prism 10.0. (D) and (E) The conidia morphology and abnormality rate of Guy11, Δ*Molst8*, and the complemented strain, bar = 10 μm. (F) Pathogenicity tests were conducted by spraying conidia onto barely leaves. The concentration of conidial suspensions for each strain was 5 × 10^4^ spores/ml, cultured for 7 days at 25°C. (G) Observation of appressorium-mediated invasive hyphal (IH) colonization on detached barley leaves for Guy11, Δ*Molst8*, and the complemented strain, bar = 10 μm. (H) The pathogenicity experiment on barley leaves was conducted using mycelial plugs from Guy11, Δ*Molst8*, and the complemented strain. Observation was carried out after 4 days of inoculation under 25°C conditions. (I) Conidial suspensions (5 × 10^4^ spores/ml) from Guy11, Δ*Molst8*, and the complemented strain were applied in the pathogenicity experiment. Observation was carried out after 4 days of inoculation under 25°C conditions. (J) The pathogenicity experiment on barley leaves using mycelial plugs from Guy11, Δ*Molst8*, and the complemented strain. (K) Pathogenicity tests were conducted by spraying conidia onto rice leaves.

### MoLst8 positive regulate the activity of TORC1

Inhibitors like rapamycin impact cell metabolism and growth by targeting the TORC1 complex [77]. MoLst8 is a TORC1 component, and we studied its relationship with MoTOR kinase under rapamycin’s influence to understand its regulatory significance [74, 78, 79]. The Guy11 and Δ*Molst8-C*, cultured with 100 ng/ml rapamycin on CM media for 8 days, exhibited significantly smaller colonies, indicating substantial growth inhibition. In contrast, the Δ*Molst8* mutant showed heightened sensitivity to rapamycin, essentially halting growth (Figure 2A and 2B). To validate the regulatory role of MoLst8 on MoTOR activity, we measured the phosphorylation level of the TORC1 activity marker Rps6 in both Guy11 and Δ*Molst8* under CM liquid media without rapamycin and with 100 ng/ml rapamycin. The results in Figure 2C revealed that the degree of phosphorylation of Rps6 in Δ*Molst8* was already weak in comparison to Guy11, indicating a significant reduction in TORC1 activity due to the absence of the *MoLST8* gene. After the addition of rapamycin, the Δ*Molst8* mutant exhibited complete loss of Rps6 phosphorylation (Figure 2D), indicating the complete inhibition of TORC1 activity in response to rapamycin. In summary, the *MoLST8* gene is a direct positive regulator of TORC1 activity.

**Figure 2.**
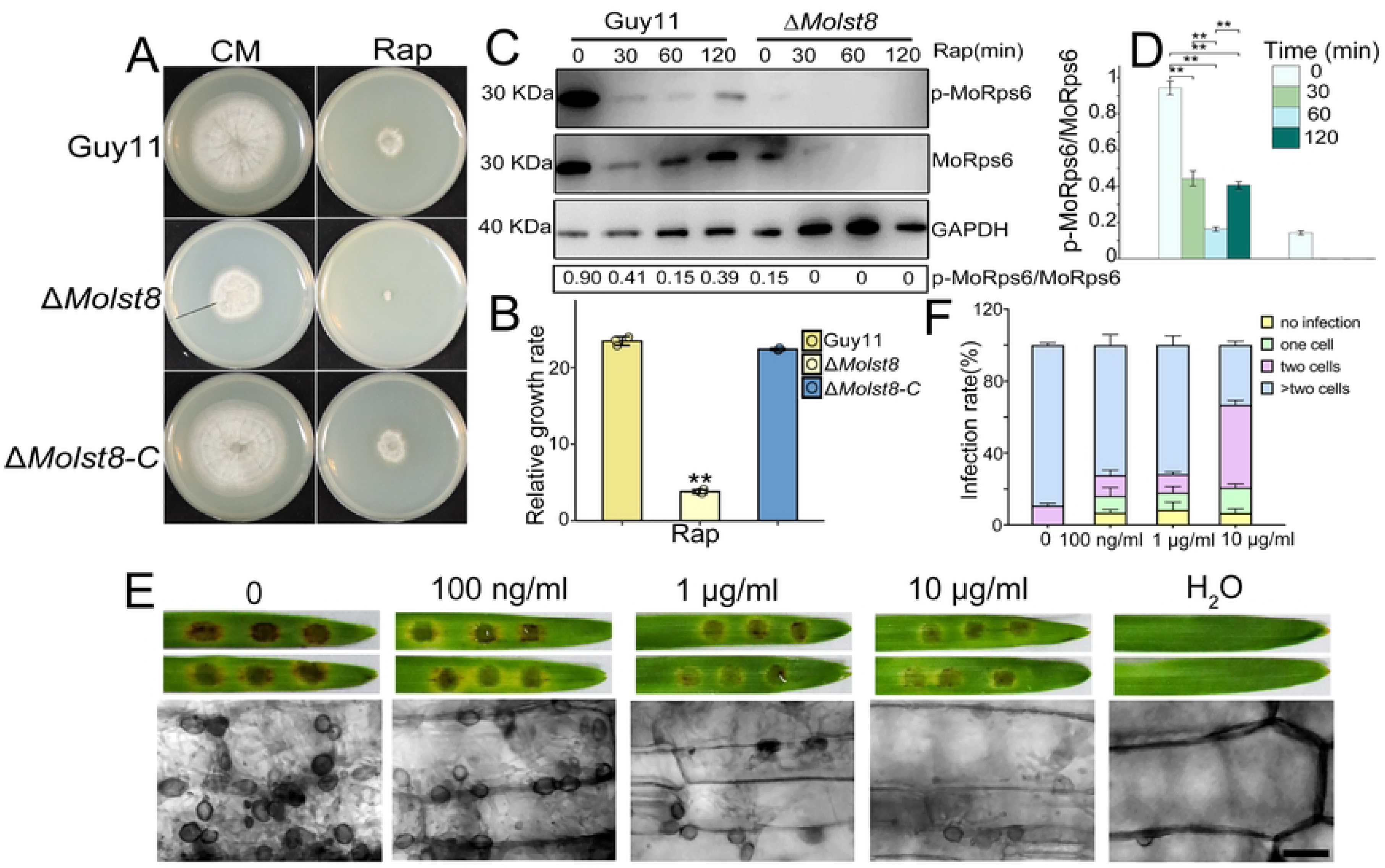
The *MoLST8* gene positively regulates TORC1 activity. (A) Guy11, Δ*Molst8*, and the complemented strain were cultured on CM supplemented with 100 ng/ml rapamycin at 25°C for 8 days. (B) Guy11, Δ*Molst8*, and the complemented strain exhibited different relative growth rates, with statistical significance denoted by double asterisks. (C) Phosphorylation analysis of MoRps6 in Guy11 and Δ*Molst8*. (D) Bar chart illustrating the phosphorylation levels of MoRps6 (** p < 0.05). (E) Detection of Guy11 conidia virulence on barley leaves under the influence of rapamycin at concentrations of 0, 100 ng/ml, 1 μg/ml, 10 μg/ml. Conidia (5 × 10^4^ spores/mL) were cultured on five-day-old barley leaves at 25°C for 4 days, the IH was observed by microscope, bar = 20 μm. (F) Quantitative statistical analysis of invasive hyphal growth for Guy11, Δ*Molst8*, and complemented strains under the influence of rapamycin at concentrations of 0, 100 ng/ml, 1 μg/ml, 10 μg/ml.

Subsequently, we examined how the major component of the rice blast fungus TORC1 complex, *MoLST8* gene, participates in regulating normal cell growth under the influence of rapamycin. Spore suspensions of wild-type Guy11 were placed onto detached barley leaves and incubated for a duration of 4 days with varying concentrations of rapamycin (0, 100 ng/ml, 1 μg/ml, 10 μg/ml). Lesions on barley leaves gradually diminished, suggesting a gradual decrease in pathogenicity even at elevated rapamycin concentrations. As rapamycin concentration increases, fewer attached appressoria invade and establish adjacent to more than two cells, while the number of those adjacent to one or two cells increases. (Figure 2E and 2F). Under rapamycin influence, attached appressoria retain the capability to produce IH and invade host cells.

### Autophagy was depressed in Δ*Molst8* mutant

Autophagy is a pivotal intracellular mechanism regulating various stages of fungal growth [14, 80, 81]. TOR regulates autophagy, shaping cellular responses to nutrition and the environment, impacting metabolism and survival [83, 84]. To explore the function of MoLst8, a major component of TOR, in regulating autophagy, we expressed GFP-MoAtg8 fusion proteins in Guy11 and Δ*Molst8* strains. Autophagic flux was assessed through fluorescence microscopy and immunoblotting. In nutrient-rich CM, GFP-MoAtg8 fluorescence predominantly appeared as punctate structures around vacuoles in the cytoplasm, with significantly fewer punctate structures in Δ*Molst8* compared to wild-type Guy11. Under amino acid and ammonium sulfate deficiency (SD-N) after 6 hours of starvation, Guy11 displayed extensive vacuole fragmentation, and GFP-MoAtg8 puncta formed larger structures inside vacuoles. These structures aggregated into a sheet-like distribution challenging to discern within and around vacuoles. As starvation progressed up to 12 hours, the sheet-like distribution of GFP-MoAtg8 fluorescence dispersed along with the fragmented vacuoles, and punctate edges became less distinct. Under starvation in synthetic media (SD-N), Δ*Molst8* exhibited a distinct response from wild-type Guy11. Most GFP-MoAtg8 fluorescence in Δ*Molst8* was within vacuoles and almost completely degraded. Punctate signals of GFP-MoAtg8 fluorescence in Δ*Molst8* significantly decreased. After 12 hours of starvation, vacuoles exhibited a fragmented distribution, and only a few punctate GFP-MoAtg8 signals were observed around vacuoles (Figure 3A-C). Subsequently, western blot analysis was conducted to assess autophagic flux by monitoring the degradation of GFP and GFP-MoAtg8 proteins. This evaluation aimed to elucidate the autophagic activity in response to nitrogen starvation conditions and provided insights into the autophagy dynamics in the fungal strains under investigation. In CM liquid media, the GFP-MoAtg8 band in Δ*Molst8* was noticeably weaker than in Guy11. Upon six hours of starvation, the band representing free GFP signal increased, while the signal corresponding to the GFP-MoAtg8 fusion protein band intensity decreased, and the degradation rate of GFP-MoAtg8 in Δ*Molst8* was slower than in Guy11 (Figure 3D and 3E). Additionally, we analyzed the endogenous lipidation of MoAtg8 using western blotting. Under nutrient restriction in SD-N, both Guy11 and Δ*Molst8* exhibited significantly higher levels of MoAtg8-PE compared to nutrient conditions. Moreover, under both CM and SD-N conditions, the Δ*Molst8* mutant showed much greater amounts of MoAtg8-PE than the Guy11 (Figure 3F).

**Figure 3.**
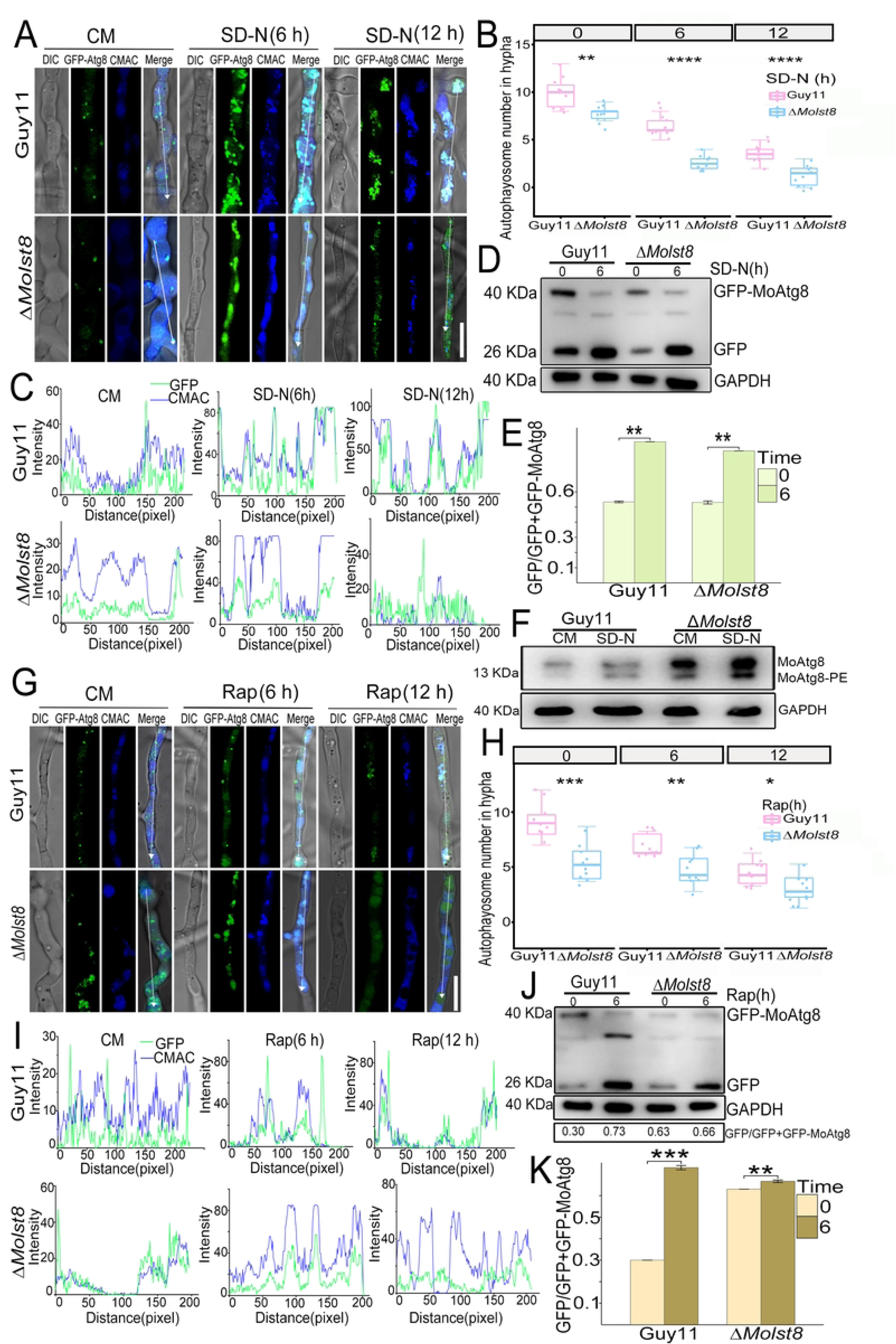
Autophagy was depressed in Δ*Molst8*. (A) GFP-MoAtg8 fluorescence assessed autophagy flux in Guy11 and Δ*Molst8* strains expressing the GFP-MoAtg8 fusion protein. After 2 days in CM medium, hyphae were shifted to SD-N for 6 and 12 hours, stained with CMAC, and examined under a fluorescence microscope, bar = 10 μm. (B) Using box plots to statistically analyze the number of autophagosomes in Guy11 and Δ*Molst8* strains under SD-N conditions. (C) ImageJ software was used to determine the colocalization of GFP-MoAtg8 and CMAC vacuolar fluorescence intensity in Guy11 and Δ*Molst8* hyphae under SD-N treatment. (D) Monitoring autophagic flux of GFP-MoAtg8 hydrolysis under SD-N conditions using western blot. (E) Bar chart showing GFP-MoAtg8 hydrolysis rate under SD-N treatment. (F) Western blotting detected the MoAtg8/MoAtg8-PE conversion in Guy11 and Δ*Molst8* in CM and SD-N medium. (G) GFP-MoAtg8 fluorescence assessed autophagy flux in Guy11 and Δ*Molst8* strains expressing the GFP-MoAtg8 fusion protein. After 2 days in CM medium, hyphae were shifted to rapamycin for 6 and 12 hours, stained with CMAC, and examined under a fluorescence microscope bar = 10 μm. (H) The number of autophagosomes in Guy11 and Δ*Molst8* strains were analyzed under rapamycin treatment for 6 and 12 hours. (I) ImageJ software was used to determine the colocalization of GFP-MoAtg8 and CMAC vacuolar fluorescence intensity in Guy11 and Δ*Molst8* hyphae under rapamycin conditions. (J) Monitoring autophagic flux of GFP-MoAtg8 hydrolysis under rapamycin conditions using western blot. (K) Bar chart showing GFP-MoAtg8 hydrolysis rate under rapamycin treatment.

Rapamycin, by inhibiting TOR activity, initiates autophagy, participating in controlling cell growth and metabolism [85]. MoLst8 is known a crucial component of TOR, studying its effect on autophagic flux in the presence of rapamycin can help researchers understand how this gene product interacts with the autophagy machinery and how it might be modulating the process. After 6 hours of rapamycin treatment, wild-type Guy11 showed intense GFP-MoAtg8 fluorescence with smaller punctate structures around vacuoles, while in Δ*Molst8*, GFP-MoAtg8 fluorescence was mostly within vacuoles and undergoing significant degradation. After 12 hours of rapamycin treatment, Δ*Molst8* exhibited more severe vacuolar degradation and GFP-MoAtg8 fluorescence degradation, with GFP-MoAtg8 fluorescence losing its punctate morphology and merging together. Meanwhile, at this time point, the GFP-MoAtg8 fluorescence signal in wild-type Guy11 sharply decreased, and GFP-MoAtg8 fluorescence appeared within vacuoles, initiating degradation (Figure 3G-I). Subsequently, autophagic flux of degraded GFP and GFP-MoAtg8 was analyzed by western blotting. After 6 hours of rapamycin treatment, Guy11 exhibited faster degradation of the fusion protein GFP-MoAtg8 and an increased rate of free GFP bands compared to Δ*Molst8*. In Δ*Molst8*, the GFP-MoAtg8 band showed no change, resulting in a slower enhancement of the free GFP band (Figure 3J and 3K).

### Identification and annotation of phosphorylated proteins in *M. oryzae*

Employing the MoMo analysis tool, which relies on the motif-x algorithm, we examined the motif features of the modification sites. For this analysis, we considered peptide sequences encompassing 6 amino acids on both sides of all detected modification sites (Fig. S4A). Based on the MoMo analysis results, a heat map was used to show the scoring of the degree of change in amino acid frequency near the modification sites (Fig. S4B). LC-MS/MS analysis identified 3220 proteins, comprising 940 differentially expressed proteins, with 614 being up-regulated and 326 being down-regulated. There were 1352 differential modification sites, wherein 933 proteins were found to be up-regulated and 419 were down-regulated (Fig. S4C). The volcano plot was utilized to represent the differentially expressed proteins between Guy11 and Δ*Molst8* mutant strain, with the top 10 labeled. The threshold was set at Log2 Fold Change > 0.5 and p-value < 0.05 (Fig. S4D). The heatmap showed significantly different phosphorylated proteins (Figure 4A).

**Figure 4.**
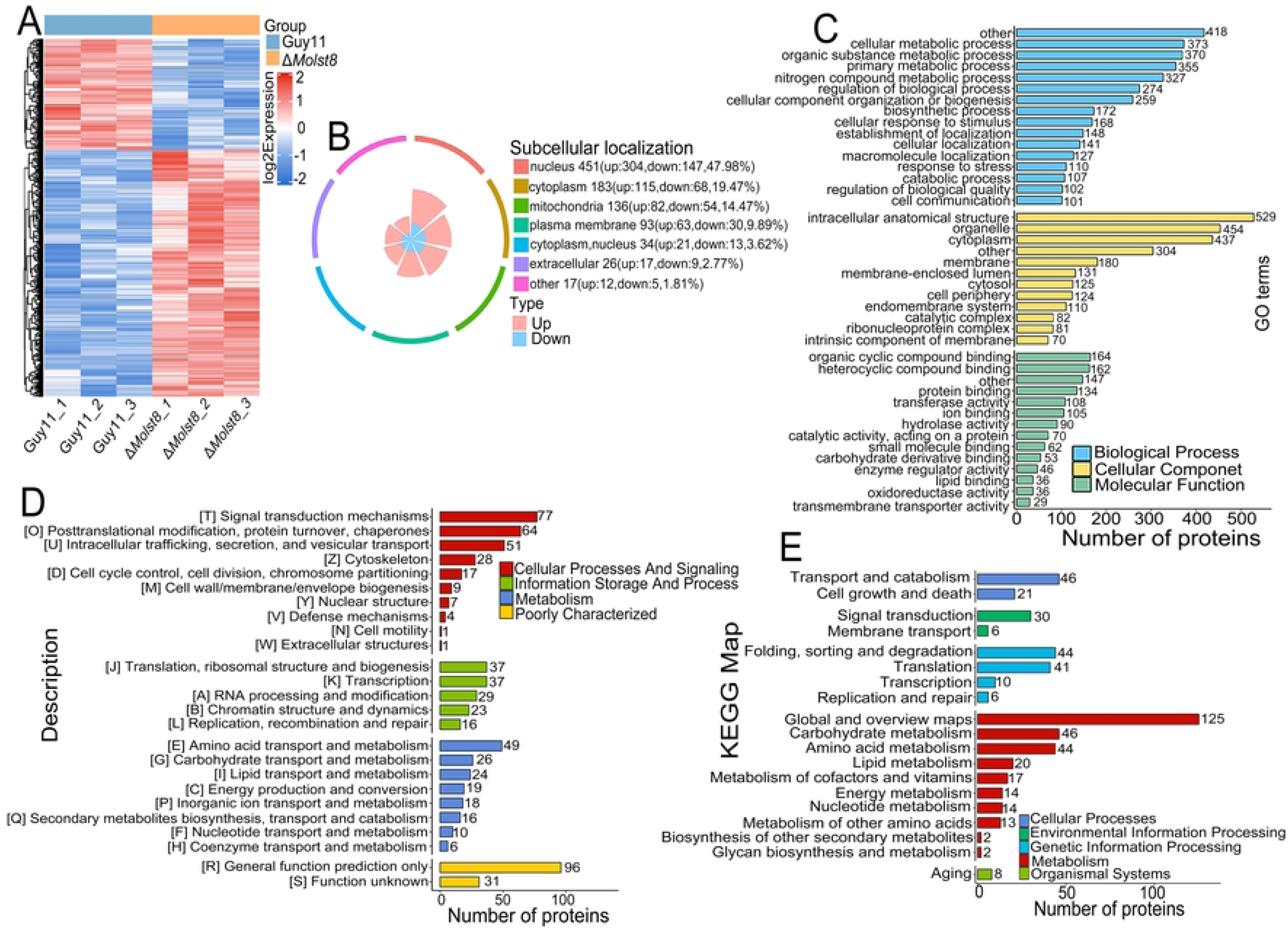
Characteristics of identified phosphorylated proteins. (A) Heatmap of differentially modified phosphorylated protein in Guy11 and Δ*Molst8* mutant. (B) Plot the subcellular localization of differentially modified phosphorylated proteins. (C) Performing gene ontology functional classification of differentially modified phosphorylated proteins based on molecular function, cellular component, and biological process. (D) Enrichment analysis of domains in differentially modified phosphorylated proteins. (E) Enrichment analysis of differentially modified phosphorylated proteins based on KEGG pathways.

To better understand the intracellular functions of phosphorylated differentially expressed proteins, subcellular localization, GO classification, COG/KOG classification, and KEGG pathway classification were performed. WoLFPSORT was used to label their subcellular localization, as shown in Figure 4B. Phosphorylated proteins were in the nucleus (47.98%), cytoplasm (19.47%), mitochondria (14.47%), plasma membrane (9.89%), cytoplasm-nucleus (3.62%), extracellular (2.77%), and others (1.81%). As shown in Figure 4C, phosphorylated proteins were enriched in cellular metabolic process (373 proteins), regulation of biological process (274 proteins), biosynthetic process (172 proteins), cellular response to stimulus (168 proteins), response to stress (110 proteins), and cell communication (101 proteins) in biological process; 180 proteins were enriched in membrane of cellular component; 36 proteins were enriched in lipid binding of molecular function. We also studied protein domains and found that they were enriched in cell wall/membrane/envelope biogenesis (9 proteins), cytoskeleton (28 proteins), signal transduction mechanisms (77 proteins) of cellular processes and signaling; 37 proteins were enriched in translation, ribosomal structure and biogenesis of information storage and processing; 24 proteins were enriched in lipid transport and metabolism of metabolism (Figure 4D). KEGG pathway analysis revealed that 19 pathways were enriched with phosphorylated proteins (Figure 4E), including cell growth and death (21 proteins), signal transduction (30 proteins), lipid metabolism (20 proteins), and membrane transport (6 proteins).

Some upregulated proteins are involved in autophagy (Atg1, Atg11, Atg2, Atg3, Atg5), lipid metabolism and plasma membrane homeostasis (Cho2, Nte1, Erg6, Psd2), replication (Tof1), and proteolysis (Pepp). Downregulated proteins participate in stress response (Nst1), transport (Sec31), signal transduction (Hse1 downregulated, Vps27 upregulated), metabolism (Hpd4 downregulated, Katg1 upregulated). Five differentially expressed proteins are involved in transcription, with three (Egd2, Nut1, Sub2) downregulated and two (Htb1, Trf1) upregulated. Five proteins are involved in RNA processing, with two (Ded1, Prp5) downregulated and three (Abd1, Spp2, Tsr3) upregulated. Four proteins are involved in translation, with one (Prt1) downregulated and three (Hcr1, Nip1, Pab1) upregulated (Table 1). In summary, phosphorylated differentially expressed proteins are involved in autophagy, PM homeostasis, signal transduction, stress response, growth and development, and metabolism.

### MoLst8 coordinate TORC2 and involve PM homeostasis

The TORC2-Ypk1 signaling module promotes the synthesis of sphingolipids by activating specific enzymes, which is crucial for cytoskeleton reorganization and cell membrane composition and function [36, 37, 86]. To reinforce the involvement of MoLst8 in lipid homeostasis, we quantified lipid levels by means of lipidomics analysis. Detailed lipid components detected in Guy11 and Δ*Molst8* samples employed in this experiment are provided in Data Sheet S1. Among the 757 quantified lipids, 351 showed significant differences between Guy11 and Δ*Molst8* (Log_2_FC >0.5 (FC, fold change) and an adjusted P-value of < 0.05), with 152 displaying negative and 199 showing positive expressions (Data Sheet S2 and Fig. S5A, B), and Figure 5A presents a heat map illustrating significant differences among 50 lipids. Enrichment results of the KEGG pathway revealed that dysregulated lipids such as Cer, DAG, LPE, PA, PC, PE, SM, TAG, etc., are involved in processes including sphingolipid metabolism, lipid metabolism, steroid biosynthesis, inositol phosphate metabolism, arachidonic acid metabolism, linolenic acid metabolism, autophagy, amino acid metabolism, phosphatidylinositol synthesis, and phosphatidylinositol signaling system (Fig. S5C and Figure 5B-D). TOR signaling inhibition in microalgae triggers triacylglycerol (TAG) accumulation. In addition, TOR inhibition also affects de novo fatty acid synthesis dependent on fatty acid synthase, which is necessary for TAG accumulation. This indicates TOR’s key role in TAG synthesis/storage regulation [91]. A study found that in Chlamydomonas, the inhibition or reduced activity of the TORC1 signaling pathway in the *lst8* mutant leads to a significant accumulation of TAG [74]. We found that many types of TAG content were also significantly increased in the *MoLST8* gene-deleted *M. oryzae*, echoing the observations made by Inmaculada Couso, et al. [74]. Among them, individual types such as LacCer d18:1/20.0, PA(14:0/18:1)+AcO, PE(0-16:0/18:3), and PS(16:0/18:0) were also significantly elevated, while other DAG, LPE, LPI, FFA, LPG, PG, PC, SM, Cer, PA and PE levels were significantly reduced (Fig. S5C, Figure 5C and 5D). These results further confirm that the deletion of *MoLST8* leads to disruption of the TOR signaling pathway, resulting in dysregulation of lipid homeostasis and autophagy.

**Figure 5.**
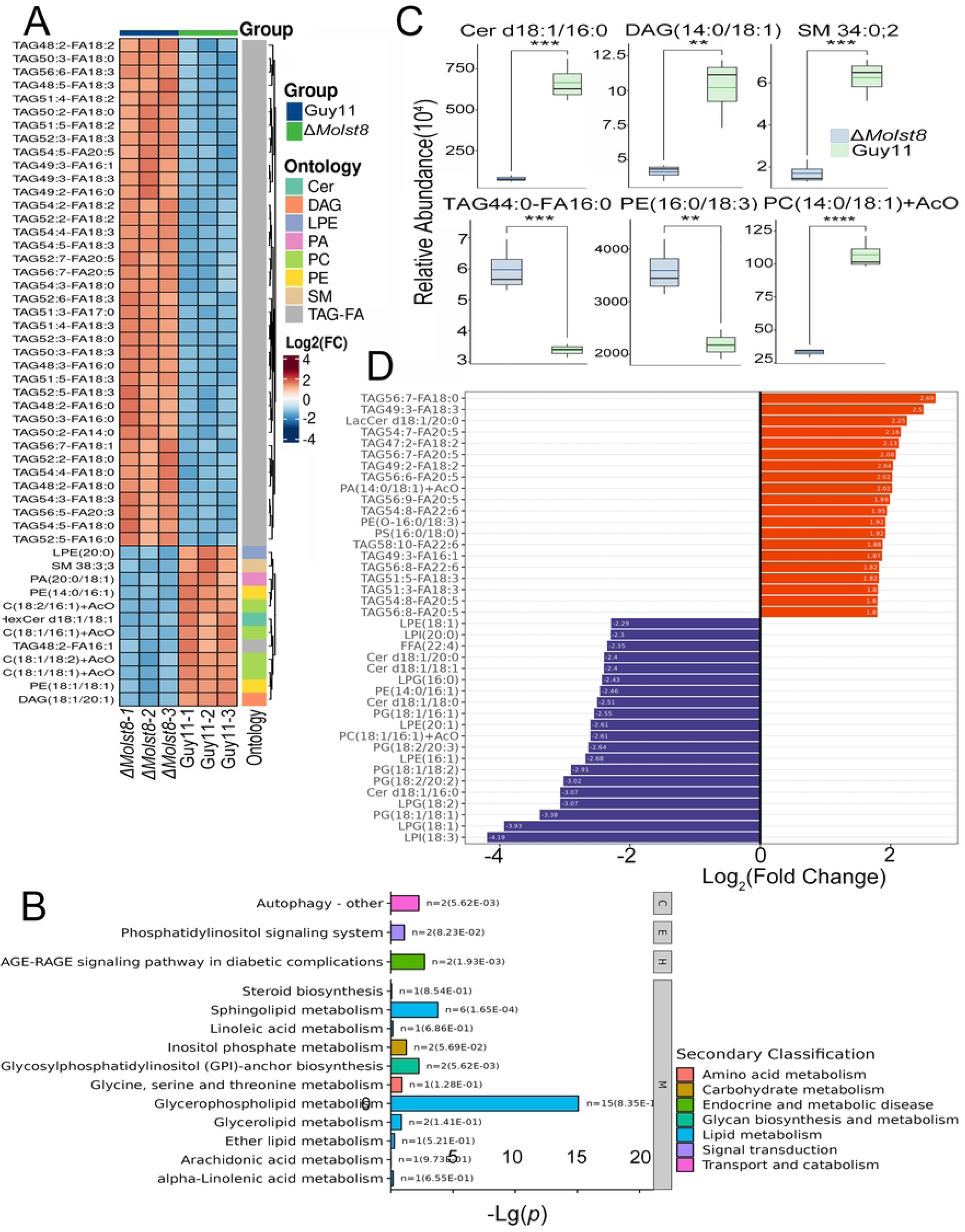
MoLst8 involved in lipid biosynthesis. (A) Heatmap shows 50 lipids with significant differences between Δ*Molst8* and Guy11. (B) KEGG analysis highlights these significant different lipids’ role in autophagy, lipid metabolism, and steroid biosynthesis. (C) Bioinformatic analysis quantifies TCA, PE, Cer, SM, DAG, sterols, and other lipids with notable variations. (D) The fold change of lipids in Δ*Molst8* mutant.

In yeast, TORC2 phosphorylates Ypk1 at T662, enhancing its kinase activity and serving as a TORC2 activity indicator [92]. In *M. oryzae*, a similar site, S619 of MoYpk1, is phosphorylated by TORC2 [24]. MoLst8 is part of TORC2, but its role in TORC2 regulation is unclear. We used western blot to observe MoYpk1 phosphorylation as a proxy for TORC2 activity, seeking a link between MoLst8 and TORC2. In CM liquid medium, the phosphorylation level of MoYpk1 in Δ*Molst8* was very low, substantially lower compared to Guy11. After treatment with 10 μM palmitoylcarnitine (PalmC), the phosphorylation status of MoYpk1 in Δ*Molst8* increased slowly but remained weak, close to zero. The phosphorylation status of Guy11 gradually decreased and then slowly recovered after 2 hours, and was much higher than that of the Δ*Molst8* mutant at any time point (Figure 6A and 6B). Since the deletion of MoLst8, a component of TORC2, leads to a significant reduction in TORC2 activity to nearly complete loss, we speculate that MoLst8 performs a key function in modulating the activity of TORC2.

**Figure 6.**
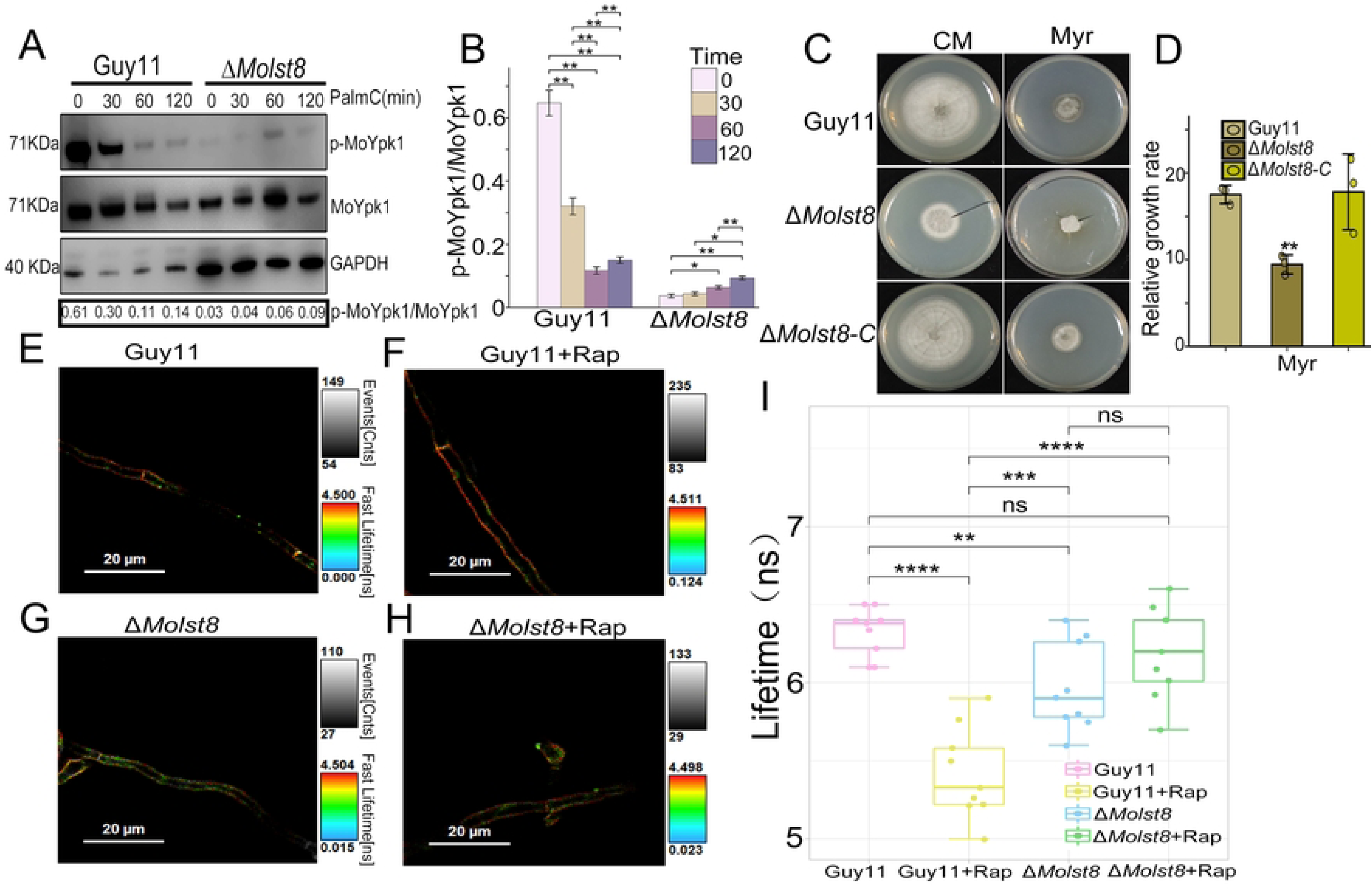
MoLst8 coordinate TORC2 and involve in PM homeostasis. (A) To study MoYpk1 phosphorylation in Guy11 and Δ*Molst8* mutants, strains were grown in CM medium for 2 days, treated with 10 μM PalmC at different time points (30, 60, 120 min), and proteins were extracted using TCA-acetone-SDS for western blot analysis on 12.5% SDS-PAGE. (B) Statistical analysis of MoYpk1 phosphorylation levels in Guy11 and Δ*Molst8* mutants using bar graphs. (C) Colony morphologies of Guy11, Δ*Molst8*, and complementary strains were examined on CM agar with 1 μM myriocin at 25°C. (D) Relative growth rates in Guy11, Δ*Molst8*, and complementary strains. (E) Fluorescence imaging of Guy11 mycelium with 2 μM Flipper-TR: longer lifetime correlates with higher PM tension. (F) Fluorescence imaging of Δ*Molst8* mycelium with 2 μM Flipper-TR: longer lifetime correlates with higher PM tension. (G) Fluorescence imaging of Guy11 mycelium treated with rapamycin and stained with 2 μM Flipper-TR: longer lifetime correlates with higher PM tension. (H) Fluorescence imaging of Δ*Molst8* mycelium treated with rapamycin and stained with 2 μM Flipper-TR: longer lifetime correlates with higher PM tension. (I) Box plots display fluorescence lifetime distributions of Guy11 and Δ*Molst8* mutant hyphae, comparing before and after rapamycin treatment. They reveal median, interquartile range, and outliers.

We are interested in how MoLst8, TORC2’s functional regulation heavily relies on it, affects plasma membrane (PM) homeostasis. Therefore, we further examined changes in sphingolipids in Guy11, Δ*Molst8* mutants, and Δ*Molst8-C* strains. Compared to the Guy11 and complementary strains, the Δ*Molst8* exhibited increased susceptibility to myriocin, a naturally occurring inhibitor of sphingolipids that interacts with and suppresses the function of serine palmitoyltransferase (Spt1) (Figure 6C and 6D). Myriocin inhibits sphingolipid synthesis to activate the TORC2-Ypk1 signaling pathway [38]. However, in our current study, we found that when the *MoLST8* gene is deleted, TORC2 activity is almost completely lost, resulting in weak Ypk1 phosphorylation levels. This prevents myriocin from effectively enhancing TORC2-Ypk1 activity and promoting sphingolipid recovery. Since sphingolipids are essential for cell growth, the Δ*Molst8* mutant exhibits further growth inhibition under the influence of myriocin. In summary, the deletion of *MoLST8* disrupts plasma membrane homeostasis, which is detrimental to cell growth.

To further understand the role of MoLst8 in maintaining normal cell morphology, we explored the membrane tension and elasticity of Guy11 strains and Δ*Molst8* mutants when exposed to rapamycin. The fluorescent membrane tension probe Flipper-TR converts fluorescence of different lifetimes into different colors for visualization, allowing precise measurement of membrane tension through fluorescence color and time [93, 94]. We used Fluorescence Lifetime Imaging Microscopy (FLIM) technology to quantify the fluorescence lifetime of wild-type and Δ*Molst8* mutants, thereby quantifying cell membrane tension [24, 48, 95]. Compared to Guy11, the filipin fluorescence lifetime of Δ*Molst8* was notably reduced. However, there was no significant difference in the change of fluorescence lifetime of Δ*Molst8* before and after rapamycin treatment, while the fluorescence lifetime of the wild-type Guy11 decreased sharply (Figure 6E-I). Our research data indicate that when the *MoLST8* gene is deleted, TORC2 activity is almost completely lost, and the inactivation of the TORC2-Ypk1 pathway will lead to a reduction in sphingolipids, which are an important component of the cell membrane and will seriously affect the lipid composition of the cell membrane, resulting in decreased membrane tension. After treating wild-type Guy11 with rapamycin, the TOR signaling pathway is blocked, affecting lipid metabolism, leading to changes in membrane fluidity, permeability, and a decrease in membrane tension. The deletion of *MoLST8* may cause the TOR complex to fail to form or lose its activity, making the TOR signaling pathway completely ineffective. In this case, rapamycin may not influence the inactive TOR complex, which may be one of the reasons why there is no significant change in cell membrane tension under the action of rapamycin in cells with deleted *MoLST8* gene.

### MoLst8 mediate TOR cross talk with MAPK pathway

We explore the interplay between TOR and MAPK signaling, emphasizing their coordinated cellular responses to environmental stresses, with a spotlight on MoLst8’s central role as a TOR component in both pathways. When Guy11, Δ*Molst8*, and Δ*Molst8-C* were grown on solid CM supplemented with 400 μg/ml CR and 0.0025% SDS, we observed that in comparison to Guy11 and Δ*Molst8-C* strains, the Δ*Molst8* mutant was extremely sensitive to CR and SDS, and its growth rate was very slow. This suggests that after the deletion of the *MoLST8* gene, the strain becomes more sensitive to CWI stress (Figure 7A-D). The mitogen-activated protein kinase mps1 and osm1 in *M. oryzae* are key to stress responses, fungicide sensitivity, and plant infection. Mps1 is central to cell wall integrity, plant infection, and stress reactions [101].

**Figure 7.**
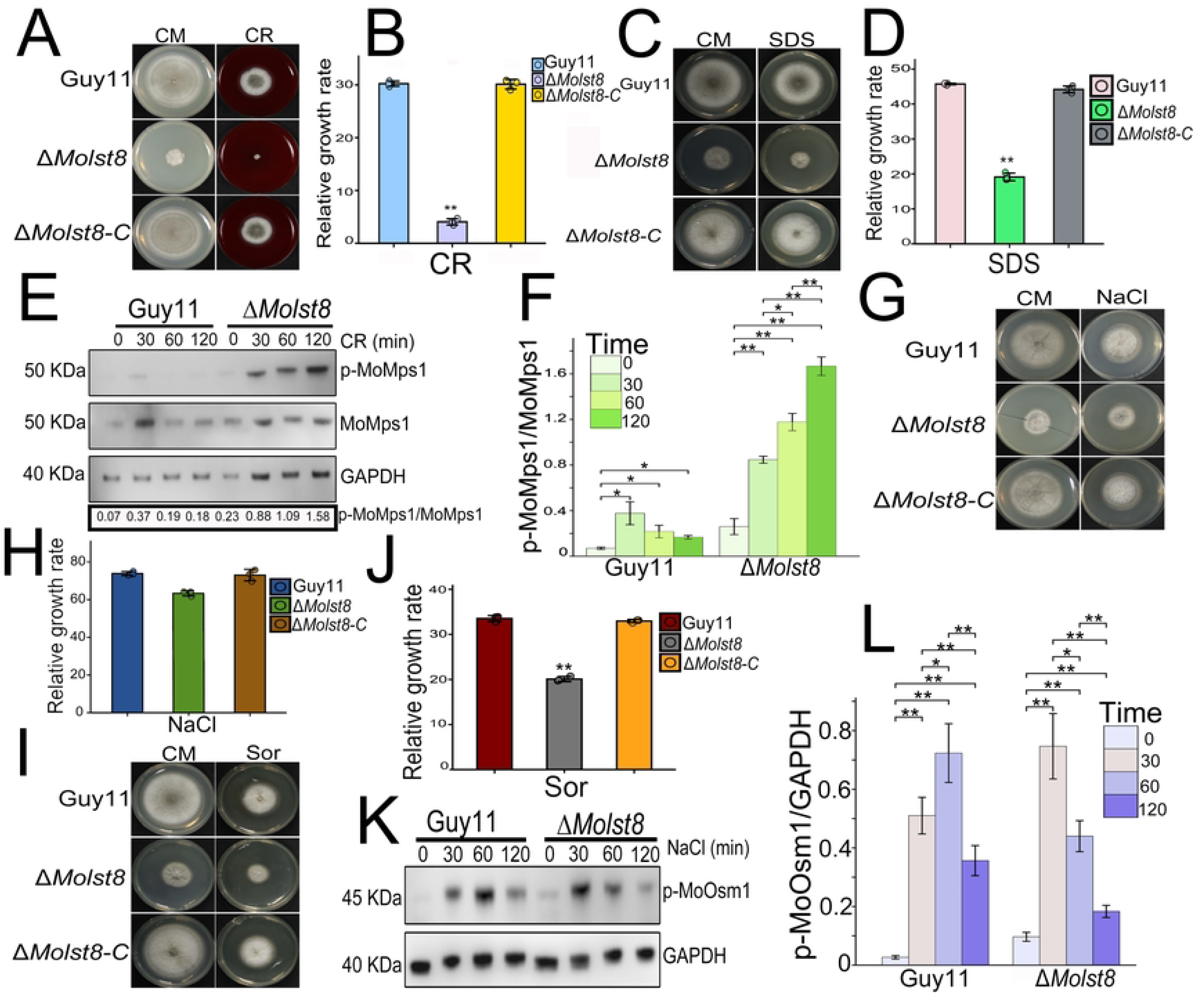
MoLst8 mediate TOR cross talk with MAPK pathway. (A) Colony morphologies of Guy11, Δ*Molst8*, and complementary strains were examined on CM agar with 400 μg/ml CR at 25°C. (B) Relative growth rates in Guy11, Δ*Molst8*, and complementary strains. (C) Colony morphologies of Guy11, Δ*Molst8*, and complementary strains were examined on CM agar with 0.0025% SDS at 25°C. (D) Relative growth rates in Guy11, Δ*Molst8*, and complementary strains. (E) To study MoMps1 phosphorylation in Guy11 and Δ*Molst8* mutants, strains were grown in CM medium for 2 days, treated with 400 μg/ml CR at different timepoints (30, 60, 120 min), and proteins were extracted using TCA-acetone-SDS for Western blot analysis on 12.5% SDS-PAGE. (F) Statistical analysis of MoMps1 phosphorylation levels in Guy11 and Δ*Molst8* mutants using bar graphs. (G) Colony morphologies of Guy11, Δ*Molst8*, and complementary strains were examined on CM agar with 0.5 M NaCl at 25°C. (H) Relative growth rates in Guy11, Δ*Molst8*, and complementary strains. (I) Colony morphologies of Guy11, Δ*Molst8*, and complementary strains were examined on CM agar with 1 M Sor at 25°C. (J) Relative growth rates in Guy11, Δ*Molst8*, and complementary strains. (K) To study MoOsm1 phosphorylation in Guy11 and Δ*Molst8* mutants, strains were grown in CM medium for 2 days, treated with 0.5 M NaCl at different time points (30, 60, 120 min). (L) Statistical analysis of MoOsm1 phosphorylation levels in Guy11 and Δ*Molst8* mutants using bar graphs.

The Mps1 mutant exhibits reduced sporulation and is deficient in cell wall penetration and infection capabilities [102]. Osm1, meanwhile, is vital for osmoregulation, with its MAPK pathway being a target for fungicides [103]. The Δ*Molst8* mutant exhibits severe defects in CWI. To further understand the role of MoLst8 in the MAPK pathway Mps1, we examined the phosphorylation level changes of Mps1 in Guy11 and Δ*Molst8* before and after Congo Red treatment. In CM liquid, the Guy11 strain shows nearly no phosphorylation of Mps1, with values close to zero, while phosphorylation of MoMps1 occurred in Δ*Molst8*. Moreover, the phosphorylation level of MoMps1 in Δ*Molst8* mutant increased with longer exposure to Congo Red. However, after CR treatment, the phosphorylation level of MoMps1 in Guy11 surged and gradually decreased after 30 minutes of Congo Red exposure (Figure 7E and 7F). These results indicate that when the *MoLST8* gene is deleted, Mps1 is activated, and the longer the duration of CWI damage stress in the Δ*Molst8* mutant, the higher the phosphorylation level of Mps1. The Mps1 in Guy11 is immediately activated after CWI damage and its activity decreases as it adapts to the adverse environment. This reveals that MoLst8 is essential for the regulation of CWI by Mps1 MAPK. We delved deeper into the function of MoLst8 in the Osm1 MAPK pathway when subjected to hyperosmotic stress. To ascertain whether Δ*Molst8* manages the phosphorylation of MoOsm1 as a response to hyperosmotic stress, Guy11, Δ*Molst8*, and Δ*Molst8-C* were grown in CM medium supplemented with either 0.5M NaCl or 1M sorbitol. In comparison to Guy11 and Δ*Molst8-C*, Δ*Molst8* demonstrated a notable decrease in growth rate (Figure 7G-J). Western blot analysis revealed that the phosphorylation level of Osm1 in both Δ*Molst8* and Guy11 rose initially and subsequently declined after NaCl treatment (Figure 7K and 7L). These data suggest that the *MoLST8* gene can respond to hyperosmotic stress in the Osm1 MAPK pathway. Our research results indicate the hypersensitivity of the Δ*Molst8* mutant to cell wall stress. The augmented phosphorylation of both Mps1 and Osm1 in the Δ*Molst8* mutant are not enough to repair its cell wall defects. The data provided here confirms that the deletion of the *MoLST8* gene leads to TOR signaling pathway blockage, reduced sphingolipids, decreased cell membrane tension, and disrupted plasma membrane homeostasis. These severely affect the regulation of cell integrity by the MAPK signaling pathway in the Δ*Molst8* mutant, making the simple activation of Mps1 and Osm1 insufficient to restore cell wall integrity. Both the TOR and MAPK pathways are major regulators of cellular stress response, growth, and development [42, 43, 100]. The deletion of *MoLST8* may disrupt the balance between TOR and MAPK. We postulate that the TOR pathway could potentially mediate environmental cues to the MAPK cascade, facilitating downstream signal transduction as a reaction to external unfavorable conditions.

## DISCUSSION

The Target of Rapamycin (TOR) pathway, consisting of TORC1 and TORC2 complexes, is essential for managing diverse cellular functions, including growth, development, autophagy, membrane tension, lipid homeostasis and stress responses [46, 47]. TORC1, composed of Tor1 with Kog1, Tco89, and Lst8, controls cell growth, nutrient absorption, and autophagy. TORC2, consisting of Tor2, Avo1, Avo2, Avo3, Bit61, and Lst8, maintains cell membrane homeostasis. Lst8, a shared subunit, links both TORC1 and TORC2 to TOR signaling [34, 35]. TOR’s activity is modulated by various proteins like MoVast1/2, MoGap1, ASD4, TIP41, and IMP1, influencing different aspects of fungal physiology and pathogenicity. Mutations in these proteins alter TOR signaling, leading to phenotypic changes [19, 24, 31, 48–50]. Our study revealed the genome of rice blast fungus contains TOR complex proteins, with MoLst8 as a significant component. MoLst8 exhibits high conservation and homology with other fungal Lst8 proteins. The Δ*Molst8* mutant differs significantly from the wild-type Guy11 in both morphology and function. This mutant grows slowly, producing fragile and abnormally shaped conidia, and displays reduced virulence on rice and barley. Its impaired ability to infect and colonize host cells suggests that MoLst8 is essential for growth, spore formation, and pathogenicity in *M. oryzae*.

Autophagy, a conserved cellular process in eukaryotes, degrades and recycles undesirable cytoplasmic components to maintain cellular homeostasis [13]. In plant pathogenic fungi, such as *M. oryzae*, autophagy proteins govern the formation of infectious structures, specifically the appressorium., which generates pressure to penetrate host cells. Disrupting autophagy impairs this function and reduces pathogenicity [14, 25, 26, 81]. Recent research has identified novel autophagy regulators, deepening our understanding of this complex mechanism, including the carbon-sensing regulator MoAbl1, the glutaminolysis modulator MoAsd4, the deacetylation protein MoSnt2, and the sterol-interacting protein MoVast1, MoVast2 regulating TOR activity to maintain lipid homeostasis, a yeast amino acid permease homologue MoGap1 and the COP9 signalosome homologue MoCsn6 [24, 48]. Autophagy, controlled by TOR signaling, is key to fungal cell balance, death, and infection. TOR adjusts cellular functions based on nutrient availability: suppressing autophagy when nutrients are plentiful but triggering cellular degradation via autophagosomes when nutrients are scarce or TOR is inhibited by rapamycin. Studying its role in plant-infecting fungi may aid in the avoidance and healing of rice blast disease [16]. MoLst8 is a crucial component of the TOR complex, and our research has found that the absence of *MoLST8* impairs TOR and autophagy. Under starvation, the Δ*Molst8* mutant strain hydrolyzes GFP-MoAtg8 slower than wild-type Guy11. With TOR-inhibiting rapamycin, Guy11 effectively adapts via autophagy, but Δ*Molst8* struggles to adapt. These results suggest that MoLst8 regulates autophagy and is essential for maintaining homeostasis and stress response in fungal cells.

Research shows that Tor activity is crucial for autophagy regulation [47, 76]. Tor kinase inhibitors like rapamycin affect cell metabolism and growth by inhibiting the TORC1 complex [77]. The *MoLST8* gene, a key TORC1 component, was studied in relation to MoTOR kinase under rapamycin’s influence. Our research demonstrates that both wild type Guy11 and a complemented strain exhibit growth inhibition with the existence of rapamycin. However, Δ*Molst8* manifests extreme sensitivity, almost completely arresting growth. *MoLST8*’s absence causes a diminution in TORC1 activity, resulting in heightened cellular sensitivity to rapamycin’s growth-inhibiting effects. This reduction in TORC1 activity is further confirmed by the decreased phosphorylation of Rps6 in the mutant, highlighting the crucial role of MoLst8 in TORC1 function. Moreover, rapamycin further suppresses TORC1 activity in the mutant, suggesting that MoLst8 mediates cellular responses to rapamycin by influencing TORC1. This establishes MoLst8 as a stimulatory regulator of TORC1. In conclusion, the *MoLST8* gene positively regulates TORC1 activity, enabling cells to adapt to rapamycin and maintain growth, metabolism, and pathogenicity.

TORC2, or Target of Rapamycin Complex 2, is a vital signaling module in fungal cells, particularly in maintaining plasma membrane (PM) homeostasis [35–37]. The PM serves as a vital barrier in fungal cells, maintaining integrity, regulating homeostasis, and defending against environmental stresses [51, 52]. Lipids have vital functions in diverse PM processes, such as material transport, endocytosis, and signal transduction [53–56]. Ist2 and Tcb proteins are involved in PM integrity and phospholipid regulation [54, 55]. Additionally, Slm proteins activate TORC2 in response to PM stress, influencing sphingolipid production [57]. MoFpk1 regulates PM homeostasis and autophagy, while MoVast2 modulates TOR activity, affecting lipid balance [48, 56]. The TORC2-Ypk1 signaling pathway regulates sphingolipid synthesis, maintaining plasma membrane homeostasis. Reduced sphingolipids or membrane changes activate this pathway, ensuring membrane homeostasis and signal transduction. Ypk1 phosphorylation by TORC2 promotes sphingolipid production, crucial for cell growth and morphology [36, 37, 86]. *MoLST8* deletion disrupts TORC2 activity, reducing Ypk1 phosphorylation and sphingolipid production. The lack of *MoLST8* gives rise to dysregulated lipid homeostasis, including elevated TAG and reduced levels of other lipid species. The Δ*Molst8* mutant exhibits heightened sensitivity to myriocin, inhibiting sphingolipid synthesis and further inhibiting growth. Additionally, *MoLST8* deletion reduces membrane tension, likely due to altered lipid composition. Rapamycin’s effect on membrane tension is minimal in the Δ*Molst8* mutant, suggesting ineffective TOR signaling. Thus, MoLst8 is crucial for TORC2 function and plasma membrane homeostasis in *M. oryzae*.

The intricate interplay between TOR and MAPK signaling pathways in fungi orchestrates cellular responses to environmental stresses, including cell wall integrity (CWI) stress and hyperosmotic conditions [42, 43, 100]. MoLst8, a central component of the TOR complex, influencing fungal growth, development, and pathogenicity. The removal of *MoLST8* causes increased susceptibility to cell wall stress, indicating its importance in maintaining cell integrity. Furthermore, *MoLST8*’s absence elevates the phosphorylation of Mps1 and Osm1 MAPKs, but this activation is insufficient to restore cell wall integrity, suggesting disrupted regulation of cell integrity by the MAPK pathway in Δ*Molst8*. These findings underscore the interconnectedness of TOR and MAPK signaling and suggest that TOR signaling may transmit environmental signals to the MAPK cascade to regulate cellular responses to adverse conditions.

The study revealed that MoLst8, a TOR complex component, plays critical roles in cell growth, membrane homeostasis, autophagy, virulence, membrane tension and cell wall stress in *M. oryzae*. MoLst8 positively regulates TORC1 activity, evident in decreased TORC1 function and rapamycin sensitivity in Δ*Molst8* mutants. It also mediates TORC2, affecting sphingolipid synthesis and membrane stability. Δ*Molst8* mutants demonstrate enhanced susceptibility to cell wall stress and augmented MAPK phosphorylation but fail to restore cell wall integrity, indicating disrupted TOR-MAPK crosstalk. Future research on MoLst8’s role in *M. oryzae* pathogenesis promises deeper insights into TOR and MAPK signaling crosstalk, potentially revealing new antifungal targets. This could lead to more effective disease control strategies, enhancing crop protection and global food security.

## Materials and methods

### Creation of mutant strains and their complements

The mutant was produced using high-throughput gene deletion approaches detailed by Lu et al. in 2014 [75]. In summary, PCR amplification with specific primers was used to obtain separate fragments of roughly 1000 bp located upstream and downstream from the *MoLST8* gene. These fragments, along with the resistance gene fragment HPH, were then seamlessly integrated into the cleaved PKO3A plasmids using a recombinase enzyme. The recombinant cassettes obtained were inserted into the Guy11 by employing the AGL1. Positive transformants were subsequently identified on CM plates containing 200 μg/mL hygromycin B and 0.5 μM 5-fluoro-2’-deoxyuridine. Further verification of the Δ*Molst8* mutant strain was carried out using PCR and southern blot analysis [24]. To assemble the complementation vector, the entire gene sequences from Guy11 were effortlessly inserted into the PKD5-GFP vector. Utilizing the ATMT approach, the vector was efficiently transferred into the Δ*Molst8* mutant strain. Verification of positive transformants was accomplished by employing a combination of molecular techniques, including western blot analysis and fluorescence microscopy.

### Phenotypic observation of strains

Phenotypic characteristics were assessed from the Guy11, Δ*Molst8* mutant, and Δ*Molst8-C* strain on CM for a duration of 8 days, followed by measurements of colony diameter and conidial production. To evaluate virulence, mycelial plugs and spore suspensions (5 × 10^4^ spores/ml) were set onto excised barley and rice leaves and incubated at 25°C for a duration of 4 days. The Guy11 conidial suspension (5 × 10^4^ spores/ml) was enriched with rapamycin at concentrations of 0, 100 ng/ml, 1 μg/ml, and 10 μg/ml and inoculated on barley leaves in vitro for 4 days, and the plaque and invasive hyphae (IH) were observed to detect infection.

### Strain stress experiments

In the drug stress experiment, wild-type Guy11, Δ*Molst8* mutants, and Δ*Molst8-C* strains were cultivated on CM medium containing 100 ng/mL of rapamycin for an eight-day duration. For the plasma membrane (PM) homeostasis test, CM was prepared with 1 μm myriocin. Cell wall stress was assayed using CM formulated within 0.0025% SDS and 400 μg/ml of Congo Red. Hyperosmotic stress was induced by preparing CM with a combination of 0.5M NaCl and 1M sorbitol. Each strain was replicated three times, and the development of colonies was noted, captured in photographs, and the diameters of each were gauged to ascertain the rate of growth inhibition.

### Monitoring autophagy flux

To monitor autophagy flux, *Agrobacterium tumefaciens* was used to transfer the PKO3A vector carrying the GFP-MoAtg8 fragment into Guy11 and Δ*Molst8* mutant strains, respectively. PCR and fluorescence microscopy were employed to verify the successful transfer. In the autophagy degradation experiment, Guy11 and Δ*Molst8* mutant strains containing GFP-MoAtg8 strains were initially propagated on CM solid medium, subsequently shifted to CM liquid medium, and cultivated at 25℃ with constant shaking at 150 rpm for a period of 48 hours. Subsequently, they were relocated to SD-N as well as CM liquid medium infused with 100 ng/ml of rapamycin, respectively, for induction periods of 6 hours and 12 hours. Autophagosomes in the hyphae were observed and counted using fluorescence confocal microscopy.

### Liquid chromatography-mass spectrometry (LC-MS) analysis

Samples were processed with a 4× volume of phenol extraction buffer containing stabilizers and inhibitors, followed by equal volume Tris-buffered phenol addition and centrifugation. The supernatant was precipitated with 0.1M ammonium acetate/methanol, cleaned with methanol and acetone, subsequently dissolved in 8M urea. After TCA addition and precipitation, the pellet was washed with pre-cooled acetone, dissolved in 200mM TEAB, and enzymatically broken down using trypsin in a 1:50 proportion overnight. Reduction with DTT and alkylation with IAA were performed, followed by peptide solubilization in a buffer solution. The peptides were then loaded onto IMAC material, washed, and phosphopeptides were eluted with 10% ammonium hydroxide. The eluents were vacuum-dried, dispersed in Liquid chromatography mobile phase A, followed by partitioning with the aid of a vanquish neo UHPLC system with a gradient mobile phase. The peptides underwent ionization within an NSI ion source and were subsequently assessed using Orbitrap Exploris 480 mass spectrometry.

### Targeted lipidomics analysis

To conduct the lipidome profiling, both the Guy11 and Δ*Molst8* were grown in CM liquid for 48 hours at a constant temperature of 25°C before being harvested and dehydrated using a freeze dryer. For every 50 mg of the dried sample, 200 μL of an aqueous methanol mixture (3:1, v:v) was incorporated and combined with 1 mL of MTBE (methyl tert-butyl ether) at a low temperature of 4°C for a 60-minute extraction process. Following this, 200 μL of water was introduced and allowed to stand for 10 minutes, the mixture was kept at room temperature. Subsequently, it was centrifuged at 4°C and 8000 rpm for 20 minutes to isolate the supernatant, which was subsequently treated with 200 μL of SDT to dissolve any precipitated protein. The proportion of lipids in each sample was ascertained by referencing the protein content. A standard volume of supernatant from each sample was extracted for vacuum drying. The dried extracts obtained were then dissolved in 100 μL of a dichloromethane/methanol mixture (1:1, v:v), and subsequently centrifuged at 4°C and 10000 rpm for 15 minutes. The supernatant obtained from this step was then subjected to LC-MS/MS analysis for lipid profiling. This comprehensive lipidomic analysis was performed by Bioprofile, adhering to instrumentation parameters outlined in previous research studies [104].

### Western blot analysis

The Guy11 and Δ*Molst8* strains were grown in CM at 25℃ with 150 rpm for 48 hours. To detect phosphorylated MoOsm1, the cultures were divided into four portions. One portion was left untreated (0 min), while the other three were transferred to CM liquid containing 0.5M NaCl and incubated at 25℃ with 150 rpm for 30, 60, and 120 minutes, respectively. To detect MoMps1 and phosphorylated MoMps1, 400μg/ml CR was added to the CM liquid medium containing the mycelia. Similarly, 100 ng/ml rapamycin was added to detect phosphorylated MoRps6 levels using anti-phosphorylated rps6 (S235/S236) antibody (Cell Signaling Technology, 2211) and anti-rps6 antibody (Abcam, ab40820) as controls. For detecting phosphorylated MoYpk1 levels, 10 uM PalmC was added, and anti-phosphorylated MoYpk1 (S619) antibody along with anti-MoYpk1 antibody (prepared by ABclonal Biotechnology Co., Ltd.) were used. Proteins were isolated employing the TCA-SDS technique [105]. To detect GFP-MoAtg8, the mycelia were incubated in SD-N media and 100 ng/ml rapamycin media for 0 and 6 hours, respectively, and then ground in liquid nitrogen. Following this, a lysis buffer composed of 50 mM Tris-HCl (pH 7.6), 150 mM NaCl, 1% Triton X-100, and 0.5 mM EDTA was used to extract the proteins. The resolved proteins on SDS-PAGE gels were then identified using GFP antibodies.

### Membrane tension assessment

The tension of the plasma membrane (PM) was assayed by employing Flipper-TR (Spirochrome, SC020), a fluorescent dye whose fluorescence lifetime varies in response to membrane curvature. Both Guy11 and Δ*MoLst8* fungal strains were cultivated in CM for 48 hours and subsequently labeled with 2 μM Flipper-TR for a duration of 15 minutes at 25 °C. Fluorescence Lifetime Imaging Microscopy (FLIM) was performed by utilizing an Olympus FV3000 microscope fitted with a PicoHarp 300 Time-Correlated Single-Photon Counting (TCSPC) module from PicoQuant. Excitation was provided by a 488 nm laser operating at a 20 MHz repetition rate, and fluorescence emission was collected through a 565–625 nm bandpass filter. The SymPhotime 64 software suite was utilized to analyze the collected lifetime data through a three-exponential decay model.

### Statistical evaluation

The fluorescence signals obtained from fluorescence and immunoblot assays were quantified using ImageJ software. The data is expressed as the average value along with its standard deviation, calculated from a minimum of three independent replicates. To determine statistical significance, with the assistance of GraphPad Prism version 10.0 software, a two-sample Student’s t-test was performed, and the significance of the findings was evaluated based on the p-value (* P< 0.05, ** P < 0.01, ** P<0.001, ****P < 0.0001).

## Acknowledgements

We extend our gratitude to Maurizio Del Poeta for his contributions to the study of sphingolipids in pathogenic fungi, as well as to Aurélien Roux and Robbie Loewith for their research on plasma membrane tension.

---

**Fig. S1.**
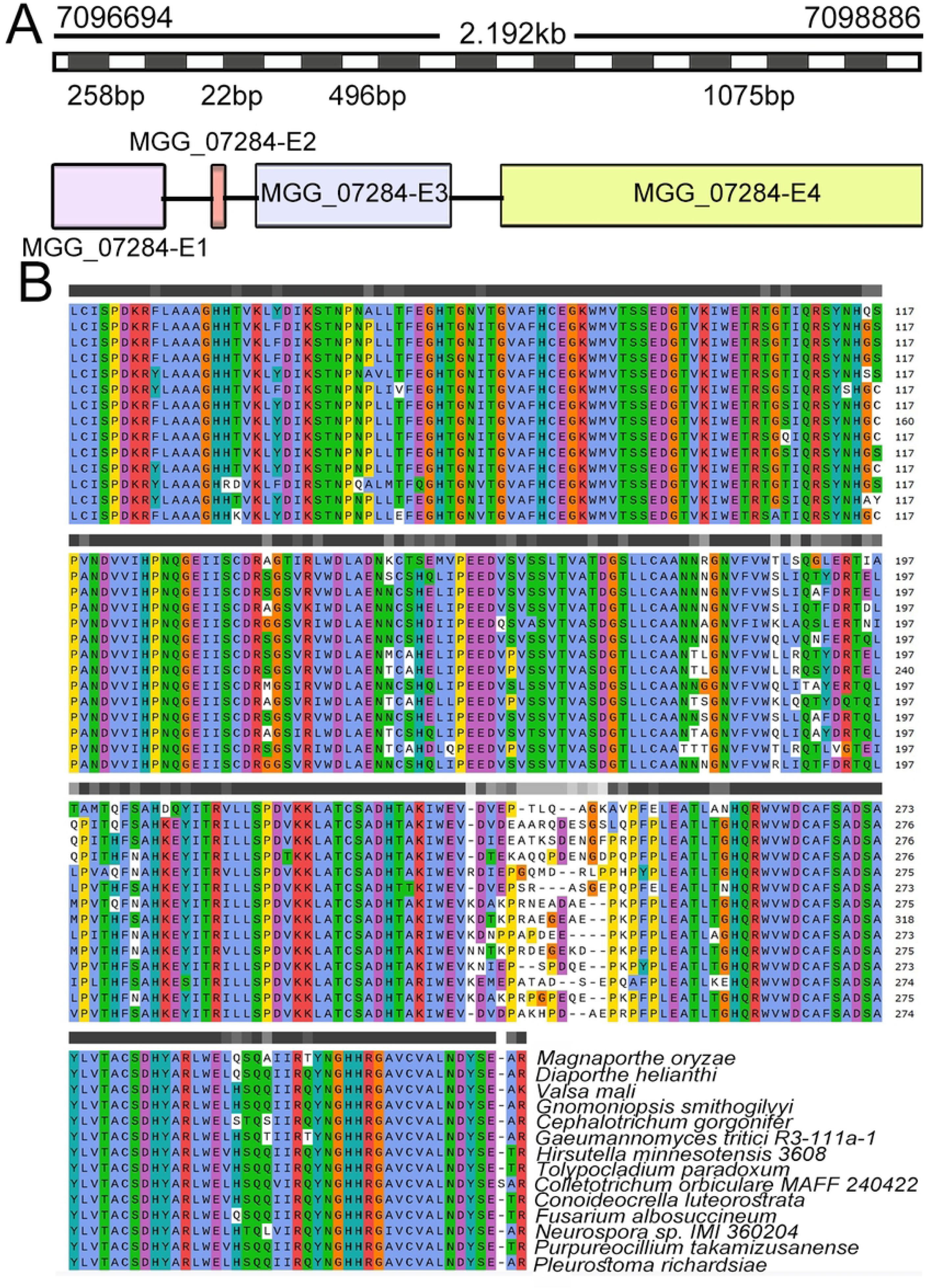
Diagram of *MoLST8* gene structure, multiple amino acid sequence alignment. (A) The gene structure of *MoLST8* in *M. oryzae*. (B) Alignment of amino acid sequences of MoLst8 from different pathogenic fungi, colors represent conserved amino acids.

**Fig. S2.**
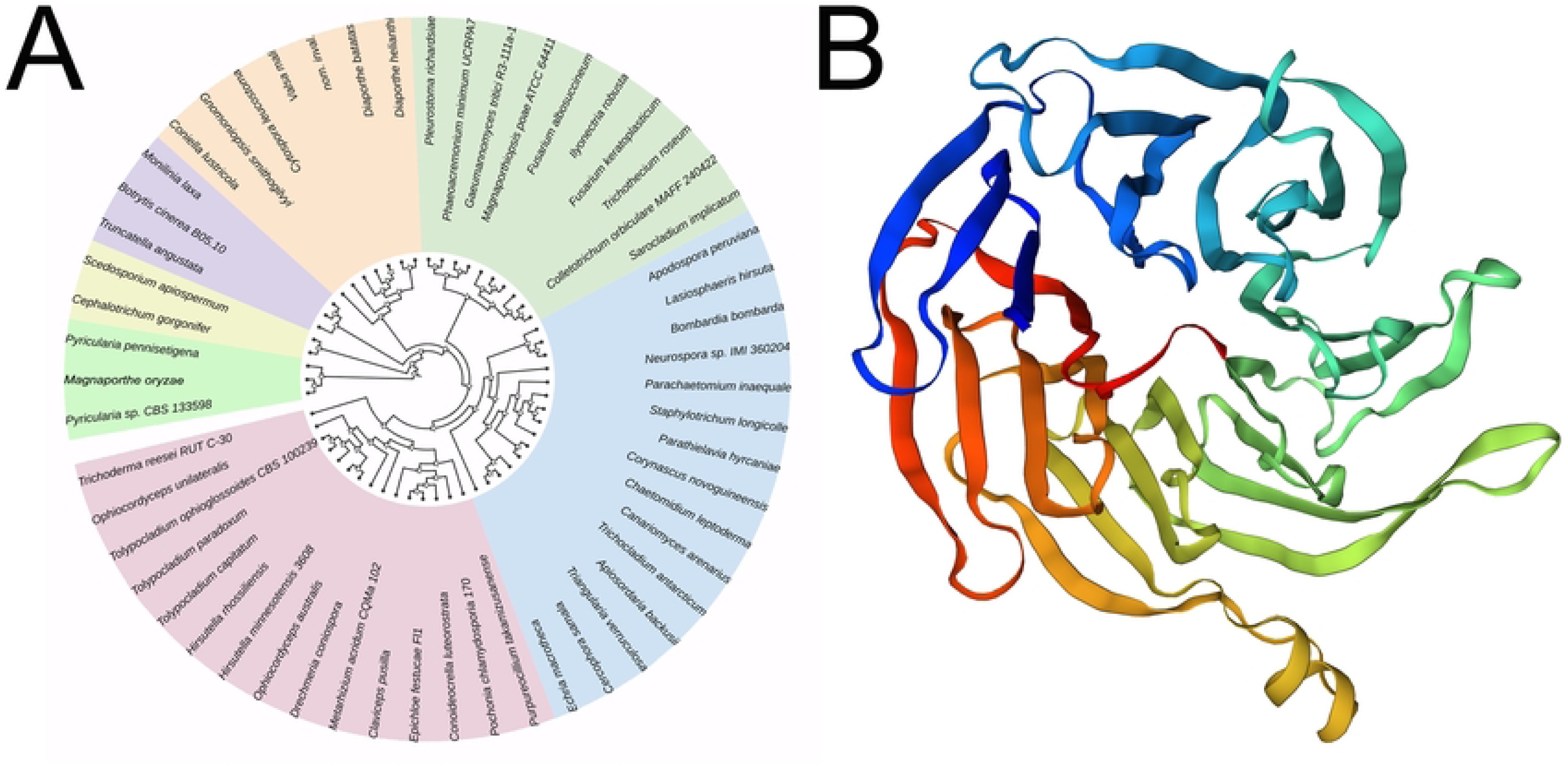
Diagram of MoLst8 structure and phylogenetic tree construction. (A) Phylogenetic tree showing the evolutionary divergence of MoLst8 with the TOR complex subunit Lst8 proteins from 13 other plant pathogenic fungi. (B) Three-dimensional structure of the MoLst8, constructed by Swiss-Model software.

**Figure S3.**
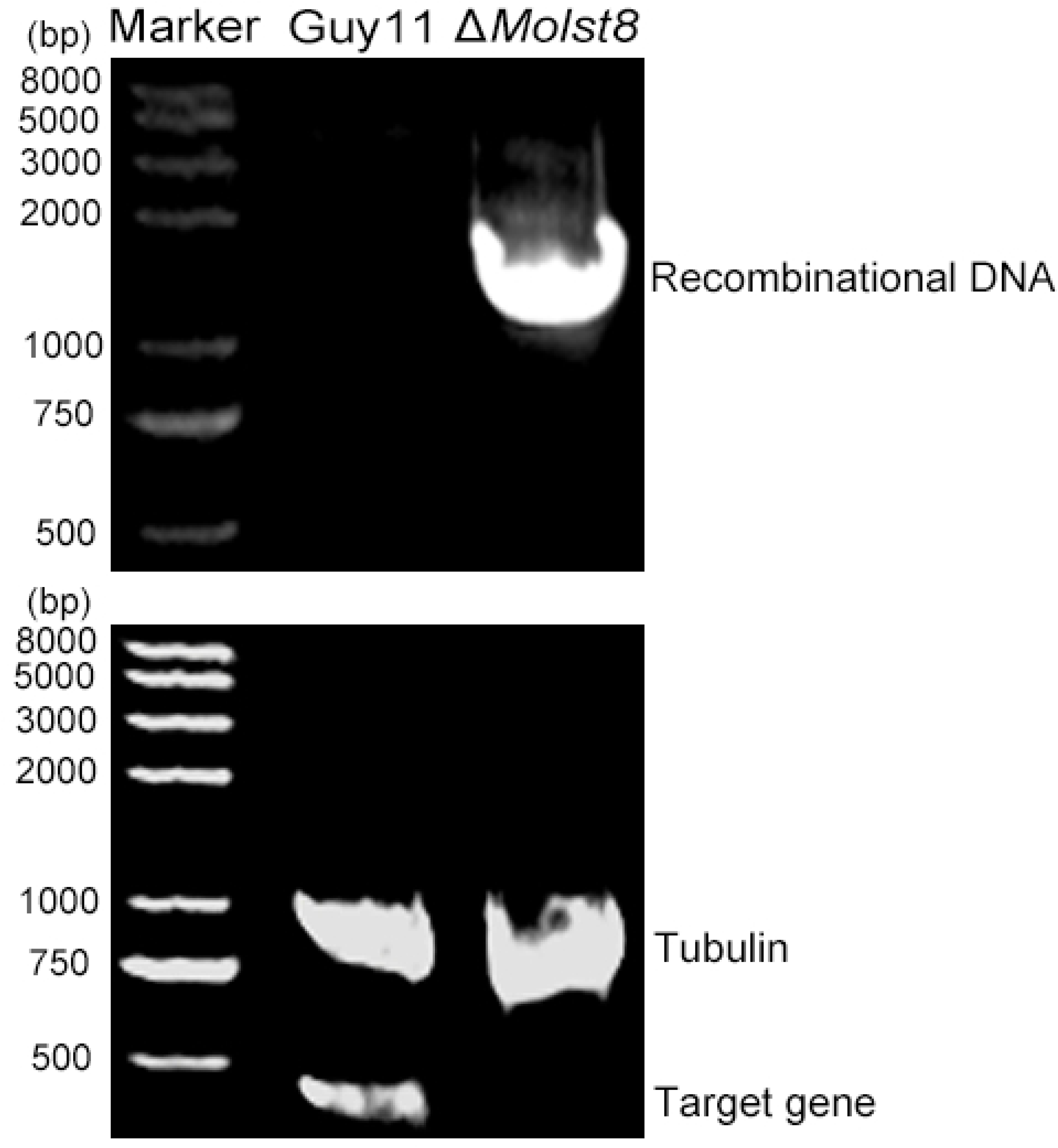
The *MoLST8* knockout verification in *M. oryzae*. PCR confirmation validated the knockout of *MoLST8*, with recombinant DNA detected at the 2000 bp position exclusively in Δ*Molst8* mutants, distinguishing them from Guy11. *MoLST8* gene presence in Guy11 contrasted with its absence in Δ*Molst8* mutants, confirming the deletion of *MoLST8* in *M. oryzae*.

**Figure S4.**
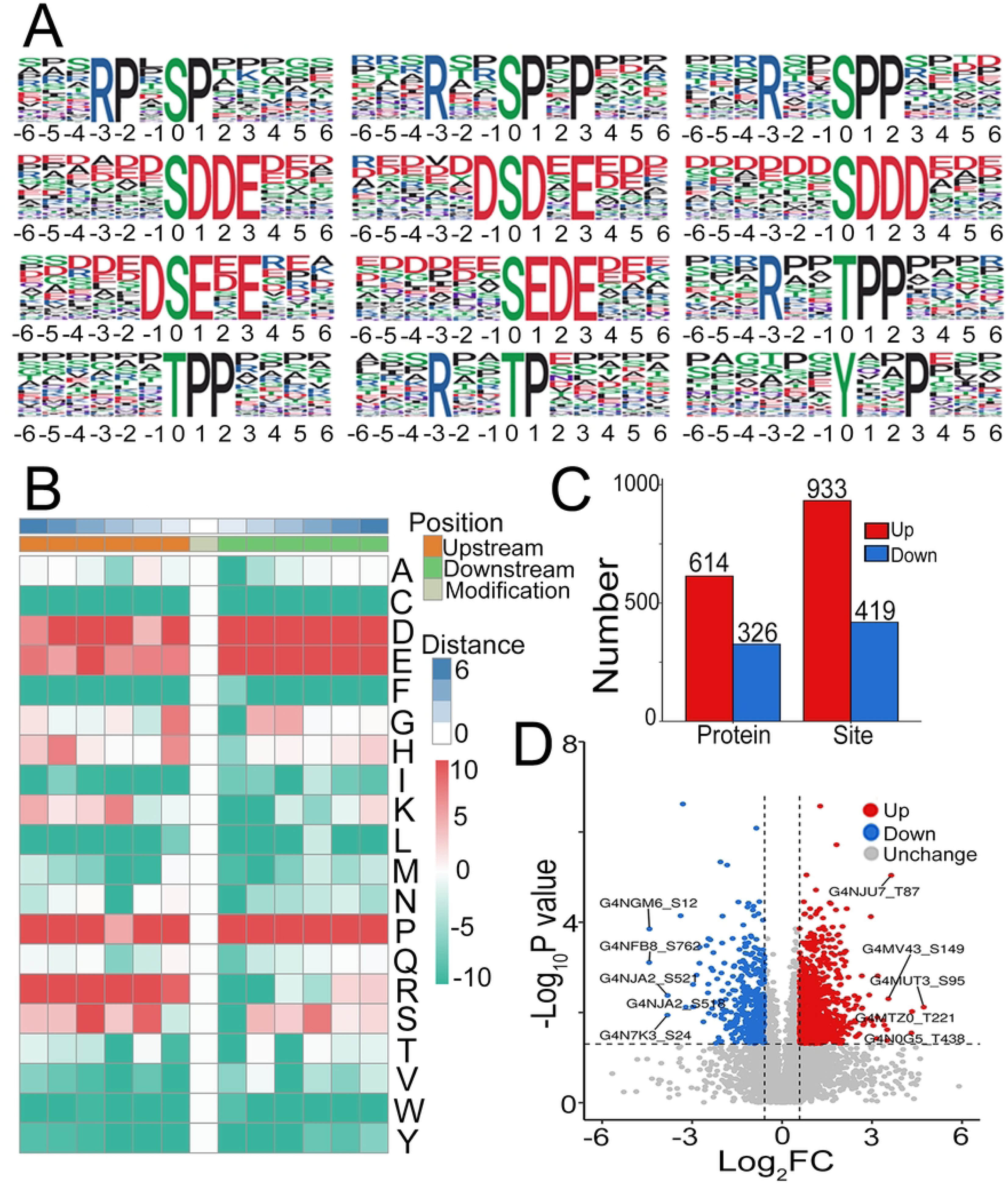
Phosphorylation motif and differentially phosphorylated expressed proteins. (A) The linear motifs enriched in phosphorylation sites detected by Motif-X software. (B) MoMo tool of motif-x algorithm analyzes the heat map of motif characteristics of phosphorylation sites. The analysis background is the peptide segment sequence composed of 6 amino acids upstream and downstream of all potential modification sites in the species. (C) Draw a histogram to visually display the distribution of differentially modified phosphorylation sites between Guy11 and Δ*Molst8* mutant. (D) Volcano plot. Red indicates up-regulated phosphorylated proteins, while green indicates down-regulated phosphorylated proteins. Five proteins with significant upregulation and downregulation differences are labeled respectively. The threshold was set at Log2 Fold Change > 0.5 and p-value < 0.05.

**Figure S5.**
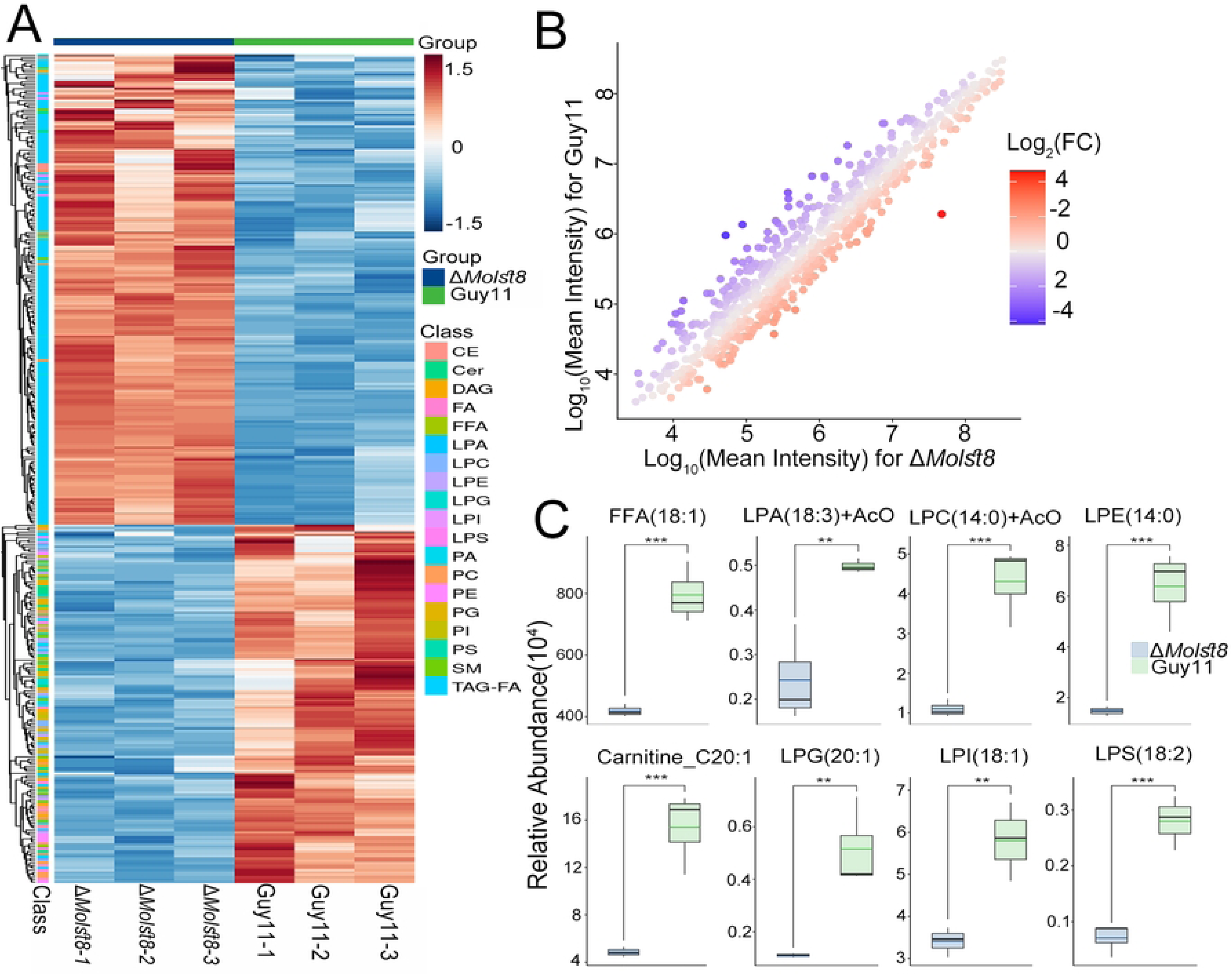
Lipidomics assessment of lipid components in Δ*Molst8*. (A) Heatmap depicting the lipid profiling differences between Δ*Molst8* and the wild-type Guy11 strain (red indicates higher lipid concentration, blue indicates lower). (B) Scatter plot visualizing the altered lipids in Δ*Molst8* compared to Guy11. (C) Bioinformatic analysis of relative abundances for lipids with significant differences.

**Table 1. Phosphorylated differentially expressed proteins in *M. oryzae***

**Data Sheet S1. Detailed lipid components detected in the wild-type Guy11 and ΔMolst8 mutant samples**

**Data Sheet S2. The significant differences of lipid components between the wild-type and Δ*Molst8* mutant**

